# Genomic reassessment of *Enterobacter* species and subspecies

**DOI:** 10.64898/2026.11.16.767154

**Authors:** Monique Clifton, Jocelyn Hume, Michael B Hall, Leah W Roberts

## Abstract

*Enterobacter* species are opportunistic pathogens whose identification is complicated by closely related lineages and changing species nomenclature. We reassessed 1,074 public genomes using average nucleotide identity (ANI), type-strain comparisons and phylogenies based on conserved markers, genome alignments and k-mers. Among 1,070 genomes included in the classification comparison, 123 changed species or subspecies membership relative to NCBI labels, 499 changed names following revisions to species boundaries or ranks, and 69 previously unnamed genomes received species assignments. The oharae, steigerwaltii, xiangfangensis and hoffmannii lineages formed an expanded *Enterobacter xiangfangensis*, separate from the lineage containing the *E. hormaechei* type. We propose five transferred combinations: *E. xiangfangensis* subsp. *oharae* comb. nov., *E. xiangfangensis* subsp. *steigerwaltii* comb. nov., *E. xiangfangensis* subsp. *hoffmannii* comb. nov., *E. roggenkampii* subsp. *quasiroggenkampii* comb. nov. and *E. chengduensis* subsp. *rongchengensis* comb. nov. We also describe the corresponding type subspecies, *E. xiangfangensis* subsp. *xiangfangensis* comb. nov., *E. roggenkampii* subsp. *roggenkampii* comb. nov. and *E. chengduensis* subsp. *chengduensis* comb. nov., and emend the descriptions of *E. xiangfangensis* and *E. hormaechei*. *E. dykesii* and *E. quasimori* are treated as later heterotypic synonyms of *E. asburiae* and *E. mori*, respectively. Within *E. xiangfangensis* subsp. *hoffmannii*, we define 25 genomic groups (genomovars) at 99.5% ANI. Conflicting tree relationships leave the position of *E. ludwigii* and the branching order of the hoffmannii-associated types unresolved. We developed Pterobacter to apply the revised classifications to assemblies, using alignment-based ANI near the species boundary and withholding assignments when the reference evidence is insufficient.

## Introduction

*Enterobacter* species exist across the One Health spectrum, occurring in soil, water, plants, and the intestinal microbiota of animals, including humans. Certain species are known as opportunistic pathogens, commonly found in bloodstream, respiratory, and other hospital-acquired infections. Their clinical importance has become more notable in recent years due to rising antimicrobial resistance (AMR), with modern *Enterobacter* populations found to carry multiple resistance determinants, including carbapenemases [1, 2, 3]. Accurate species identification is therefore essential for early outbreak detection and AMR surveillance. However, distinguishing closely related *Enterobacter* species remains challenging, and species misidentification continues to be reported [4, 5].

Much of this difficulty is concentrated in the *Enterobacter cloacae* complex, a group of closely related species within the genus. In clinical settings, routine biochemical identification and matrix-assisted laser desorption ionisation–time of flight mass spectrometry (MALDI-TOF MS) cannot consistently distinguish its members [2, 5]. Single gene sequencing, including 16S rRNA and the more variable *hsp60* gene, is also limited in its ability to precisely discriminate species as it only captures a small part of the available genomic diversity [6, 7].

While whole-genome comparisons have reshaped these classifications, a single agreed taxonomic assignment framework for *Enterobacter* has not yet been achieved. This is most evident when exploring the history of *Enterobacter hormaechei*. Following its original description, the species was divided into subsp. *hormaechei*, subsp. *oharae* and subsp. *steigerwaltii* [7, 8]. In 2018, Sutton *et al.* added subsp. *hoffmannii* and transferred *E. xiangfangensis*, originally described as a separate species, to *E. hormaechei* subsp. *xiangfangensis* [9, 10]. In 2020, Wu *et al.* instead supported separate *E. hormaechei*, *E. hoffmannii* and *E. xiangfangensis* species, placing the oharae and steigerwaltii lineages with xiangfangensis [5]. In 2025, De Maayer *et al.* recovered three corresponding genomic groups and proposed subdivisions within xiangfangensis [3]. More recent descriptions of *E. intestinihominis* and *E. hoffmannii* subsp. *nototheniae* add further named taxa to this part of the genus [11, 12, 13, 14]. While these studies have progressively refined *Enterobacter* taxonomy, incorporating successive revisions into reference databases can lead to significant discrepancies. Consequently, the same genomic lineage may be identified and reported under different names, complicating outbreak detection, comparisons and interpretation. A coherent genus-wide taxonomic framework is therefore needed to bring these advances into alignment and support consistent identification, particularly for clinically important species.

Average nucleotide identity (ANI) provides a quantitative basis for comparing genomes. Values near 95–96% are widely used to delineate bacterial species, often alongside digital DNA–DNA hybridisation (dDDH) [15, 16]. Subspecies boundaries are less standardised; Sutton *et al.,* for example, used a 98% ANI threshold to examine subdivisions within *Enterobacter* species [10]. Phylogenetic analyses complement these comparisons by inferring evolutionary relationships from sequence variation within established statistical frameworks, allowing assessment of support for genomic lineages and their relationships to the type strains that anchor taxonomic names. Comparing different tree building approaches, such as trees inferred from conserved markers, genome alignments or shared k-mers, can allow for the identification of evolutionary relationships that are consistently recovered and those that need closer examination, particularly where recombination or uneven sampling complicates the history.

Earlier genome-based studies established much of the current framework for *Enterobacter* classification [3, 5, 10]. Here, we reassess species and subspecies assignments across the genus using a collection of public complete assemblies supplemented with type-strain genomes. We compare a phylogeny inferred from conserved bacterial markers with two distance-based phylogenies, derived from recombination-aware genome alignments and k-mer similarity. We examine disagreements between these trees alongside ANI and alignment coverage to revise the affected groups and propose the corresponding nomenclatural changes. We then compare the resulting assignments with NCBI labels and Kraken2 classifications, distinguishing genomes assigned to a different group from groups whose boundaries or rank have changed. Finally, we develop and evaluate Pterobacter, an assembly classification tool that applies this taxonomy and uses alignment-based ANI for cases near the species boundary. For closely related genomes, it also reports finer genomic groups where the reference comparisons support an unambiguous match.

## Methods

### Genome collection and sequence preparation

A total of 1,079 genome assemblies from were downloaded from NCBI, specifically those submitted as *Enterobacter* and designated complete (accessed 5^th^ February 2026). After removing duplicate submissions, poor quality sequences, and uncertain taxonomic identities, 1,074 study assemblies were included for analysis (see Supplement S01). A further 16 draft type-strain genomes were added, which were absent as the initial collection was restricted to complete assemblies. These 16 draft assemblies were added to 17 other type-strains already in the collection, to give a total of 33 distinct type-strains. Two outgroups, *Cronobacter sakazakii* (GCA_000982825.1) and *Klebsiella aerogenes* (GCA_019048125.1), were included for phylogenetic analysis. Figure 1 shows the dataset selection; Tables S01 and S02 and Supplement S01 identify the assemblies, type strains, outgroups, exclusions and dataset overlaps.

**Figure 1.**
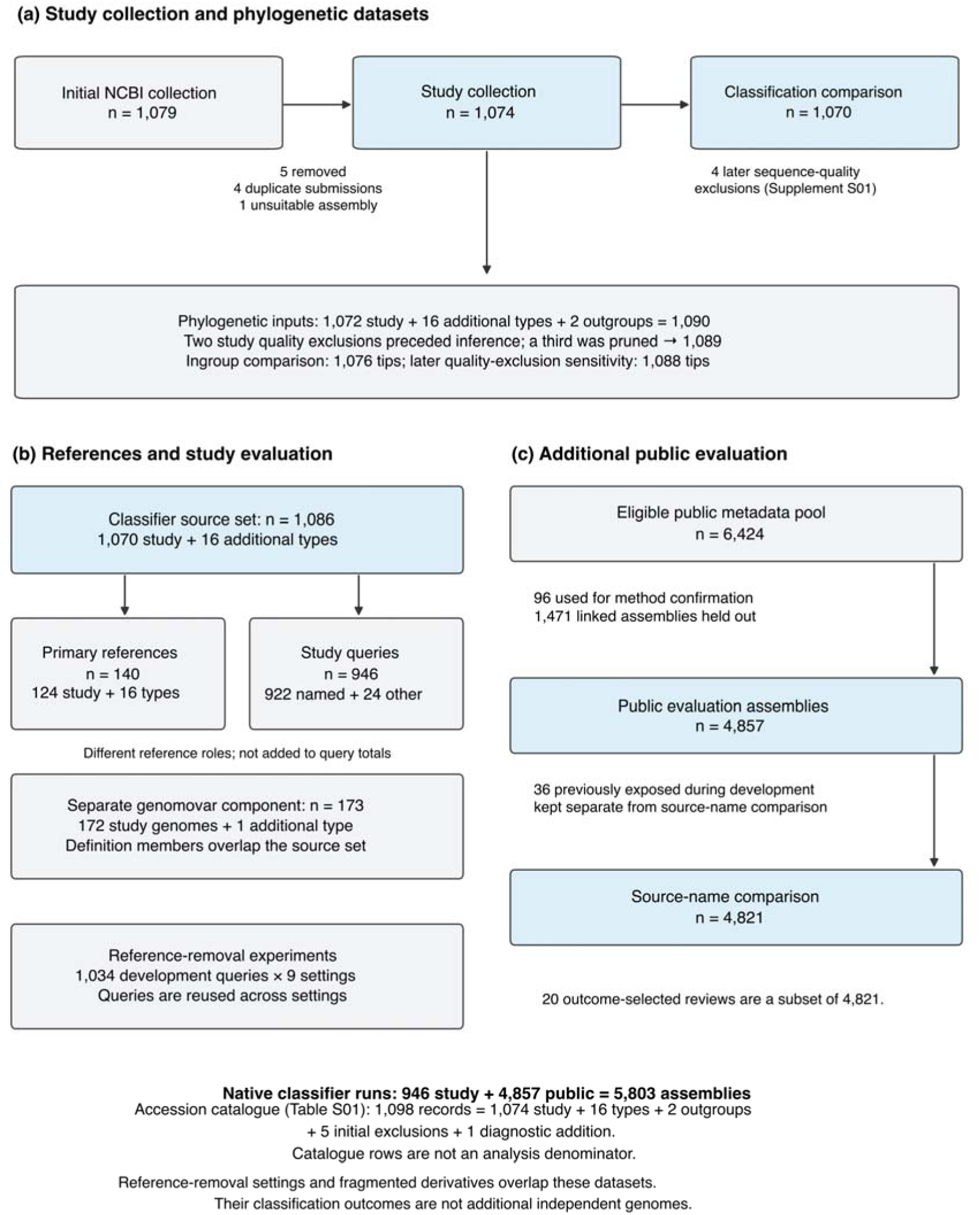
Assembly selection and the relationship between datasets. (a) Selection of the study collection and phylogenetic datasets. Four duplicate submissions and one unsuitable assembly were removed initially; four later sequence-quality exclusions account for the final 1,070-genome comparison. (b) Adding 16 type genomes gives 1,086 genomes for reference selection: 140 references (124 study and 16 additional types) and 946 non-reference study queries. (c) Additional public assemblies and classifier evaluation. The 6,424 eligible metadata records were divided into 96 method-confirmation assemblies, 1,471 related assemblies held out and 4,857 evaluation assemblies. Of the latter, 36 previously used in development were excluded from the 4,821-genome source-name comparison. The 1,098-row accession catalogue has a separate accounting purpose: 1,074 study genomes, 16 additional types, two outgroups, five initial exclusions and one diagnostic addition. Repeated reference-removal experiments and fragmented derivatives are detailed in Supplement S08; their outcomes are not added as independent genomes.

Taxonomic comparisons used chromosome sequences to limit the contribution of plasmids. All contigs were retained for draft types, whose chromosome and plasmid sequences could not be distinguished reliably. Four later quality exclusions (including one after phylogenetic inference) left 1,070 study assemblies for the final comparison of classifications. Sequence preparation and these exclusions are described in Supplement S01; tests of alternative sequence inputs are described in Supplement S05.

### Screening genomic relationships with ANI

We compared the study chromosome sets with the 33 type-strain genomes, all-vs-all, using skani 0.3.1 [17]. ANI ≥95% and aligned fraction (AF) ≥80% for each genome identified candidate species relationships for further examination. Within-group and between-group comparisons were then considered alongside the phylogenies. Selected boundaries were further examined with alignment-based ANI (ANIm) using pyANI and NUCmer, and with published dDDH measurements (Supplement S02).

### Phylogenetic inference

Three phylogenetic approaches were evaluated: conserved bacterial markers, recombination-aware genome distances, and k-mer distances. The principal comparison contained 1,089 genomes: 1,071 study assemblies (after three sequence-quality exclusions), 16 additional draft type-strain genomes, and two outgroups (Supplement S01). GTDB-Tk 2.7.2 with GTDB release R232 identified and aligned 120 bacterial marker proteins (BAC120) [18, 19]. The alignments were filtered to retain positions containing an unambiguous standard amino acid in at least 80% of genomes, and a marker tree was inferred with FastTree 2.2.0 using WAG and gamma rescaling [20].

For the genome-alignment approach, Verticall 0.4.2 estimated distances from sequence classified as vertically inherited, and FastME 2.1.6.3 inferred the tree by balanced minimum evolution [21, 22]. The k-mer approach used Mashtree 1.4.6, Mash 2.3 and quicktree 2.5, with k = 21 and 10,000 hashes per genome [23, 24]. Because changing the sketch seed changed the inferred relationships, we summarised 1,000 seed replicates as a majority-rule consensus, assigning each retained branch its frequency across replicate trees and its mean length across replicates containing that branch. Branch frequencies therefore measure stability to sketch sampling rather than bootstrap support.

The agreement and conflict among trees on identical tip sets were compared, accounting for short branches and weakly resolved relationships. Supplement S03 gives sequence preparation, branch filters and sensitivity analyses; Figure 2 shows the type-strain relationships.

**Figure 2.**
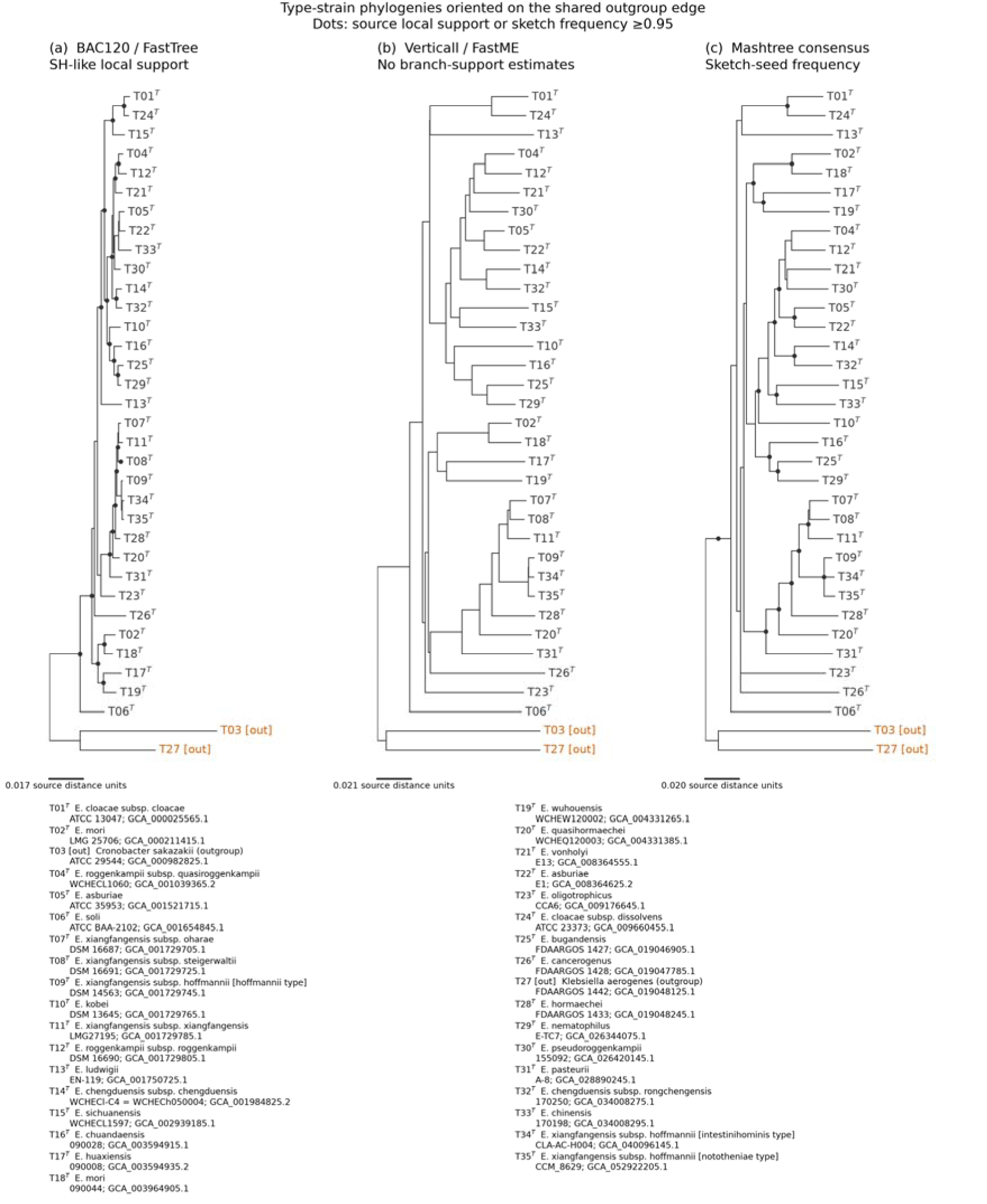
Relationships among type-strain genomes in the three phylogenetic analyses. The 35 tips represent 33 type-strain accessions and two outgroups, pruned from the 1,089-tip trees. Labels show the proposed classification. Superscript T marks the selected genomes of historical type strains; the figure key gives their deposited strain labels and genome accessions, and Table S03 maps historical and proposed names. The three hoffmannii-associated types retain separate historical-name aliases. Branch lengths and support values are those of the original trees. Dots indicate Shimodaira–Hasegawa-like (SH-like) local support ≥0.95 in the marker tree and sketch-seed frequency ≥0.95 in Mashtree; these measures have different interpretations. FastME has no branch-support estimates. For display, each 35-tip projection is rooted on the existing edge separating the two outgroups, which are marked [out]. Reorientation preserves all unrooted splits, pairwise path lengths and edge-support values; it does not establish a root for the full collection (Supplement S03). The later exclusion was assessed in a separate 1,088-tip sensitivity analysis.

### Investigation of conflicting relationships between phylogenetic approaches

Where tree topology was inconsistent between different phylogenetic methods, we examined individual core genes to determine which relationships they support, and whether that support varies across the genome. We investigated the position of *E. ludwigii*, the relationship between *E. sichuanensis* and *E. chinensis*, and the branching order of the *E. hoffmannii*, *E. intestinihominis* and *E. hoffmannii* subsp. *nototheniae* types. Core genes were identified using Pyrodigal 3.7.1 and Panaroo 1.8.0 and were aligned using MAFFT 7.526 [25, 26, 27]. Individual core-gene trees built with IQ-TREE 3.1.1 [28] were inspected to determine support for different tree topologies. We also examined where the supporting genes occurred along the genome to determine whether genes favouring the same relationship were concentrated in particular genomic regions (Supplement S04).

### Taxonomic interpretation and genomic subdivisions

Species and subspecies assignments were determined based on genomic similarity to type strains, within-group and between-group ANI, alignment coverage and phylogenetic relationships. Where genomic comparisons indicated close relationships between groups with different species names, we assessed whether they should remain separate species or be combined, with subspecies retained where supported. We report evidence conflicting with each proposed treatment and identify cases that remain unresolved (Supplement S06). Proposed names were checked against LPSN, original descriptions, Validation Lists and the International Code of Nomenclature of Prokaryotes, 2025 Revision [29, 30] (Supplement S11).

Within the proposed *E. xiangfangensis* subsp. *hoffmannii*, we defined genomic subdivisions (genomovars) by requiring every pair in a group to meet 99.5% ANI. This threshold lies near the midpoint of the reported within-species ANI gap of 99.2–99.8% [31]. The groups have no formal nomenclatural rank; clustering, threshold sensitivity and comparisons with other subdivisions are described in Supplement S07.

### Comparison with NCBI and Kraken classifications

Established taxonomic assignments were compared with NCBI names for the 1,070 eligible study assemblies, while distinguishing change in biological group from a change in the name or rank of the same group. For example, a broad historical *E. hormaechei* label could become *E. xiangfangensis* under the proposed boundaries without implying that the earlier identification was erroneous. A subspecies name referring to a different lineage instead counted as a change in group membership. Newly resolved, retained and unresolved assignments were recorded (Table 1; Supplement S01).

**Table 1.**
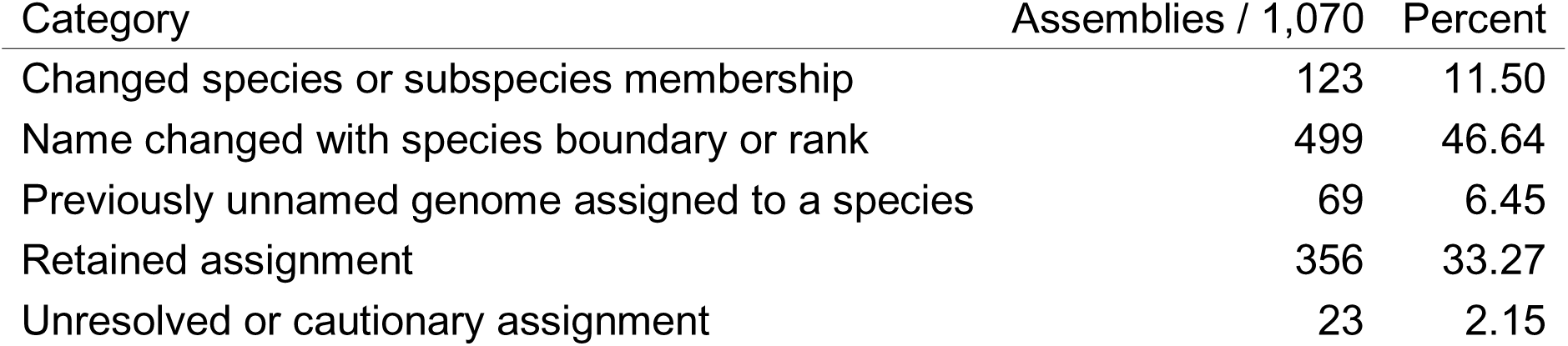
Changes from NCBI labels to the revised classification.

| Category | Assemblies / 1,070 | Percent |
|---|---|---|
| Changed species or subspecies membership | 123 | 11.50 |
| Name changed with species boundary or rank | 499 | 46.64 |
| Previously unnamed genome assigned to a species | 69 | 6.45 |
| Retained assignment | 356 | 33.27 |
| Unresolved or cautionary assignment | 23 | 2.15 |

We then compared these classifications with Kraken2 2.1.6 using the Standard database dated 26 February 2026, at confidence 0 and 0.1 [32]. Each assembly was classified as its original contigs and as artificial fragments cut into lengths of up to 150 bp, to examine how shorter inputs affected the names returned. Both the Kraken classification of the sequences and the naming of the Kraken classifications were evaluated (Supplement S09).

### Genome identification with Pterobacter

We developed Pterobacter to apply the revised taxonomy to whole genome assemblies, including plasmids and draft contigs. A panel of 140 references were chosen to represent the type strains and sampled diversity; a separate component contains the genomes defining the genomovars. The tool first compares a query with the reference panel using skani and applies alignment-based ANI near the species boundary. It reports species, subspecies and genomovar assignments only where the corresponding identity, coverage and competing-reference checks are satisfied. Reference selection, assignment rules and handling of uncertain matches are described in Supplement S08, with software versions in Supplement S10.

### Evaluation datasets and reference-removal experiments

We evaluated whether Pterobacter reproduced the study assignments and how its classifications compared with labels for additional public genomes. The first comparison used the 946 study assemblies absent from the primary reference panel. The second used 4,857 additional, non-complete NCBI assemblies, extending evaluation beyond the complete assemblies used for the taxonomic study. After classification, we also reviewed twenty selected disagreements with public names to distinguish conflicting database identifications from unsupported tool assignments. These outcome-selected cases were analysed separately (Figure 1; Supplement S08).

To test behaviour when reference coverage was incomplete, we removed the query’s close relatives or all representatives of its expected species, subspecies or genomovar. These experiments asked whether the classifier withheld an assignment or incorrectly matched a remaining group. Fragmented derivatives tested the effect of shorter assembled sequences on genomovar matching. The same genomes were reused across settings, so classification outcomes are not independent biological samples. Supplement S08 gives the exclusions, dataset overlaps and assessment of withheld or incomplete results.

### Computational assistance

OpenAI Codex and Anthropic Claude assisted with code development, computational workflows, evidence checks and manuscript drafting and revision during September–October 2026. The authors directed this work and are responsible for reviewing the outputs and the final text and scientific conclusions.

## Results

### Genome datasets

We reassessed species and subspecies assignments in public complete *Enterobacter* assemblies from NCBI RefSeq/GenBank, using type-strain genomes to anchor the names. Of 1,074 study assemblies, 1,070 remained eligible for the classification comparison after quality assessment. Additional draft assemblies were included to represent taxa absent from the complete assembly collection; Figure 1 outlines the phylogenetic and classifier-evaluation datasets in this collection (Table S01; Supplement S01).

### ANI-led reassessment of species and subspecies

We first used ANI and AF to identify relationships between study genomes and named types. Comparisons within and between candidate groups, together with the three phylogenies, then informed species and subspecies boundaries. The largest revision concerns the lineages historically placed within *E. hormaechei* (Figure S1).

We propose an expanded *E. xiangfangensis* containing 611 genomes: 438 associated with the oharae, steigerwaltii and xiangfangensis types, and a 173-genome lineage containing the hoffmannii, intestinihominis and nototheniae types, including one additional reference genome. All nine ANIm comparisons between types from these two sets gave 95.919–96.020% identity, with minimum AF ≥84.781%. Population ANI and the three phylogenies also support their treatment as one species (Figures S1 and S2; Table 2).

**Table 2.** Genomic evidence and adopted rank interpretations.

| Comparison | Type/panel evidence | Adopted interpretation and retained limitation |
|---|---|---|
| hoffmannii lineage / oharae–steigerwaltii–xiangfangensis lineages | Nine type ANIm comparisons 95.919–96.020%; minimum AF ≥84.781%; 74,445/75,774 skani pairs pass 95/80 | One xiangfangensis species with differentiated subspecies |
| oharae–steigerwaltii–xiangfangensis lineages / hormaechei type lineage | Chromosome skani 0/438 pass; whole-type 7/438; chromosome ANIm 274/438 identity passes, no AF pass | Separate hormaechei species; estimator and reference-scope dependence |
| cloacae/dissolvens | Type ANIm 95.318%; minimum AF 83.18%; 200/205 panel ANIm pairs pass | One species, two existing subspecies; five comparisons involving strain CZ862 AF were exceptions |
| mori/quasimori | Type skani 96.23%; minimum AF 80.59% | One mori species; boundary flag retained |
| roggenkampii/quasiroggenkampii | Type ANIm 95.802%; minimum AF ≥88.72%; published dDDH 64.8% | One species with proposed subspecies; contradictory dDDH remains |
| asburiae/dykesii | Type ANIm 96.336%; minimum AF 83.94%; middle-genome affinity and topology differ | One broad species; outside pair unassigned |
| bugandensis/nematophilus | Type skani 95.46%; ANIm 95.581%; coverage passes | Separate cores despite overlap; boundary pair unassigned; masked trees exploratory |
| sichuanensis/chinensis | Type ANIm 92.484–92.485%; all AF >83.22% | Separate species; one chinensis genome and unresolved alternate-type provenance |
| chengduensis/rongchengensis | Type ANIm 95.733%; minimum AF 78.88%; all six cross skani pairs fail AF; published dDDH 64.4% | One species with proposed subspecies; AF/dDDH conflict preserved |
| pseudoroggenkampii/quasiroggenkampii | Type ANIm 94.992%; 3/30 panel ANIm pairs pass | Separate cores; ECC-249 unassigned |

ANI, average nucleotide identity; ANIm, ANI calculated from nucleotide alignments with MUMmer; AF, aligned fraction; dDDH, digital DNA–DNA hybridisation. Here, “95/80” denotes ANI ≥95% with AF ≥80% for both genomes. Pairwise observations share genomes. Type measurements and sampled comparisons inform the proposed rank interpretations; published dDDH values were not remeasured. For rongchengensis, the numerical source is Table 1 of the original description, whose prose transposes the comparison strains.

Within this species, the oharae, steigerwaltii and xiangfangensis subdivisions are retained as subspecies. ANI among their 437 assigned genomes was 98.18–100% within subspecies and 96.77–98.11% between subspecies (Figure S3). The hoffmannii, intestinihominis and nototheniae types shared 99.09–99.12% skani ANI and are included together in the proposed *E. xiangfangensis* subsp. *hoffmannii* (Figure S1; Supplement S04). Population analyses identified several internal groups but did not recover a coherent three-way division corresponding to the hoffmannii, intestinihominis and nototheniae names, supporting their treatment within one subspecies while describing finer genomic variation through informal genomovars.

*E. hormaechei* remains a separate species, comprising the thirteen-genome lineage containing its type strain (Figures S1 and S2). Sequence selection effects on its type-genome comparisons are described in Supplement S05.

We treat *E. cloacae* and *E. dissolvens* as one species with two subspecies. Although close to the species threshold, their type genomes shared approximately 95.17% BLAST-based ANI (ANIb), and all 205 cross-lineage ANIm comparisons exceeded 95%. All three trees recovered the two lineages together (Figure S4). Five coverage exceptions are retained in Supplement S06.

We treat *E. dykesii* and *E. asburiae* as one species under the earlier name *E. asburiae*, and *E. quasimori* and *E. mori* as one species under the earlier name *E. mori*. Accordingly, *E. dykesii* and *E. quasimori* are proposed as later heterotypic synonyms (type skani ANI 96.12 and 96.23%; Figures S5 and S6). Heterotypic synonyms are names based on different types considered to belong to the same taxon; ‘later’ identifies the subsequently published name [30]. We also propose *E. quasiroggenkampii* and *E. rongchengensis* as subspecies of *E. roggenkampii* and *E. chengduensis*. Type ANIm of 95.802 and 95.733%, respectively, and the grouping of their lineages in the phylogenies support these treatments, despite published dDDH values of 64.8 and 64.4% [5, 33]. Table 2 and Supplement S06 hold the conflicting measurements and coverage qualifications (Figures S7 and S8).

ANI overlap did not always support merging named species. *E. bugandensis* and *E. nematophilus* retained distinct groups: within-group skani ANI was 97.69–100%, compared with 94.69–95.73% between them, and local trees retained their separation. We retain both species and leave two boundary genomes unassigned (Figure S9; Supplement S06). Other uncertain genomes receive species-only assignments, remain unnamed or remain species-unassigned (Table S01).

### Phylogenetic agreement and unresolved relationships

To distinguish consistent groupings from relationships sensitive to the sequence evidence, we compared trees based on conserved marker proteins, recombination-aware genome distances and k-mer distances. All three recovered several groups supporting the treatments above, but disagreed on some relationships, including the neighbours of *E. ludwigii* and *E. sichuanensis* (Figure 2). We examined these conflicts using individual core genes and their genomic positions (Supplements S03–S04).

For *E. ludwigii*, core genes supported both a relationship with *E. cloacae* and one with the *E. asburiae*/*E. roggenkampii*/*E. kobei* group. Genes supporting each relationship occurred in blocks along the genome. As such, the analysis retains *E. ludwigii* as a distinct species, but does not confidently resolve which other species relationship is closest (Figures S4 and S5; Supplement S04).

For *E. sichuanensis*, the Verticall and Mashtree phylogenies placed *E. chinensis* closest, whereas the marker tree associated it with *E. cloacae*. Core-gene comparisons supported the sichuanensis–chinensis relationship in 65.7–78.8% of comparisons, while conserved markers were concentrated in regions supporting the alternative. We retain *E. sichuanensis* and *E. chinensis* as separate species: their type ANIm of 92.484–92.485% is below the species threshold despite their close phylogenetic relationship (Figures S4 and S6; Supplement S04).

### Genomovars within the hoffmannii subspecies

For finer identification within the proposed *E. xiangfangensis* subsp. *hoffmannii*, we required every pair within a genomic group to have ANI ≥99.5% [31]. The 173 genomes formed 25 informal genomovars: ten with multiple genomes and fifteen singletons; the largest contained 92 genomes (Figure S10). Eleven pairs of genomes assigned to different genomovars also met this threshold. Their groups remained separate because merging them would introduce other pairs below 99.5%. These subdivisions have no formal nomenclatural rank (Supplement S07).

### Changes to public genome classifications

We compared the revised assignments with NCBI names, distinguishing a change in biological group from a change in name or rank. Among 1,070 eligible assemblies, 123 changed species or subspecies membership: 95 between species and 28 within species. A further 499 changed names following revised boundaries or ranks, and 69 gained a named-species assignment. Compatible assignments were retained for 356 genomes; 23 remained unresolved or cautionary (Table 1; Table S01).

### Classification with a general-purpose Kraken database

We compared Kraken2 Standard database classifications with the revised taxonomy to examine how the reported names would differ. At default confidence, contig calls carrying explicit conflicting subspecies names represented 37.073% of all input bases; mapped species conflicts accounted for another 0.387%. A prominent example was *E. hormaechei* subsp. *xiangfangensis* returned for steigerwaltii, hoffmannii and oharae sequences (Figure 3). With shorter input sequences or higher confidence thresholds, Kraken assigned fewer sequences to species or subspecies, instead returning broader taxonomic assignments or leaving them unclassified. Supplement S09 gives the full comparison, unmapped names and database-overlap assessment.

**Figure 3.**
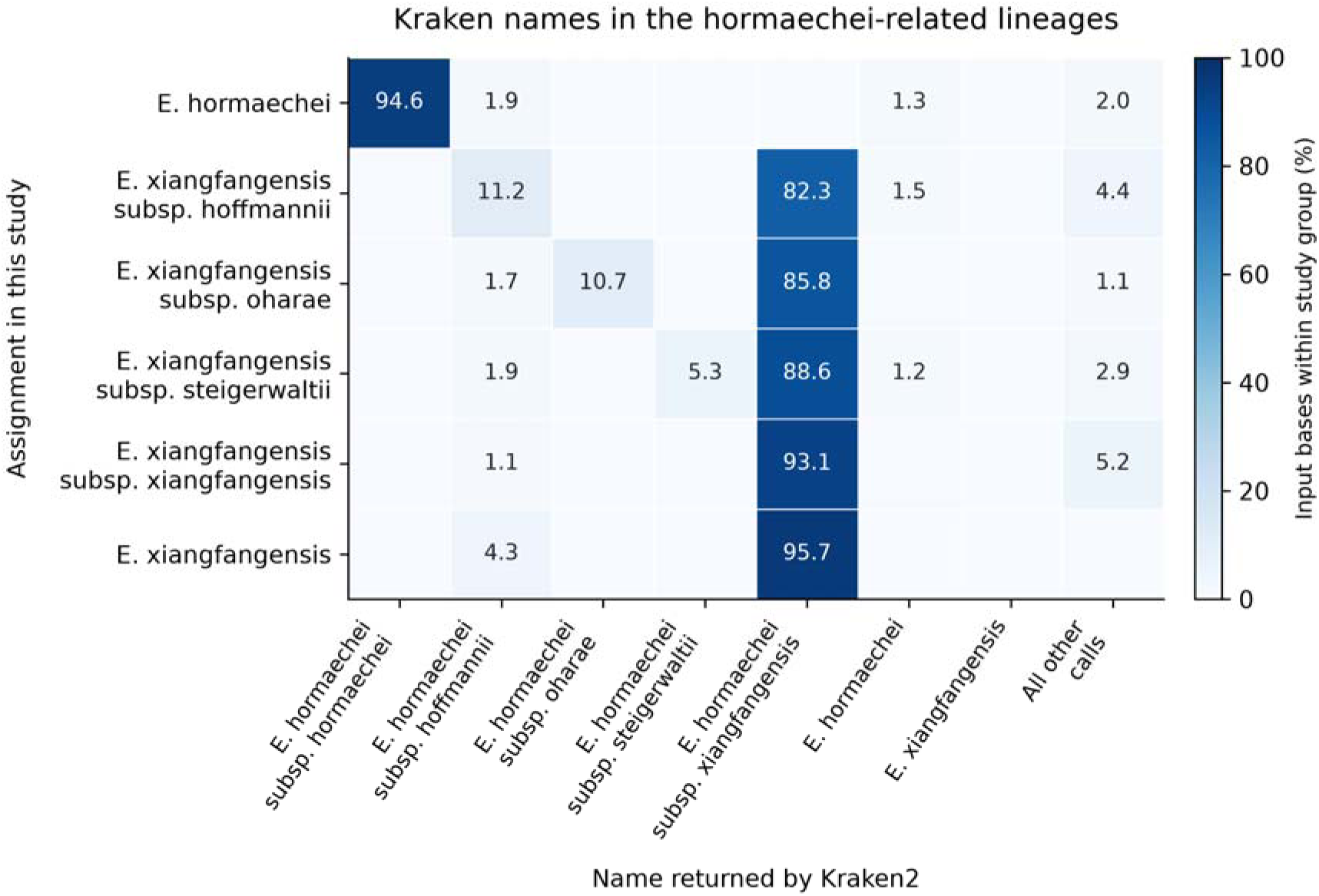
Kraken names compared with the revised assignments in the hormaechei-related lineages. Rows give the study assignment and columns the name returned by Kraken2 Standard 20260226 for native contigs at default confidence. Strain-level calls are grouped under their recorded species or subspecies ancestor. Each cell is the percentage of input bases within that row carrying the column’s sequence-level call; each sequence is weighted by its full length. All other calls are retained in the final column. The panel shows the principal subspecies disagreement, including calls to *E. hormaechei* subsp. *xiangfangensis* for three other lineages. Figure S13 gives all 32 study-assignment categories, including unnamed and contextual groups; Figure S14 shows the input-length and confidence effects on resolution. Different name strings need not imply different biological groups, and cell values are not genome-level error rates.

### Assembly identification with Pterobacter

We developed Pterobacter to identify genome assemblies according to the revised taxonomy. It compares each query with 140 references without placing it in a tree. We tested 946 study genomes excluded from this panel and 4,857 additional, non-complete public assemblies. The skani-based route assigned species to 5,671/5,803 assemblies (97.73%). It reproduced 914/922 named study assignments, withholding eight. Among public genomes with comparable names, 4,277 assignments agreed, 56 differed and 54 were withheld (Figure 4; Table 3).

**Figure 4.**
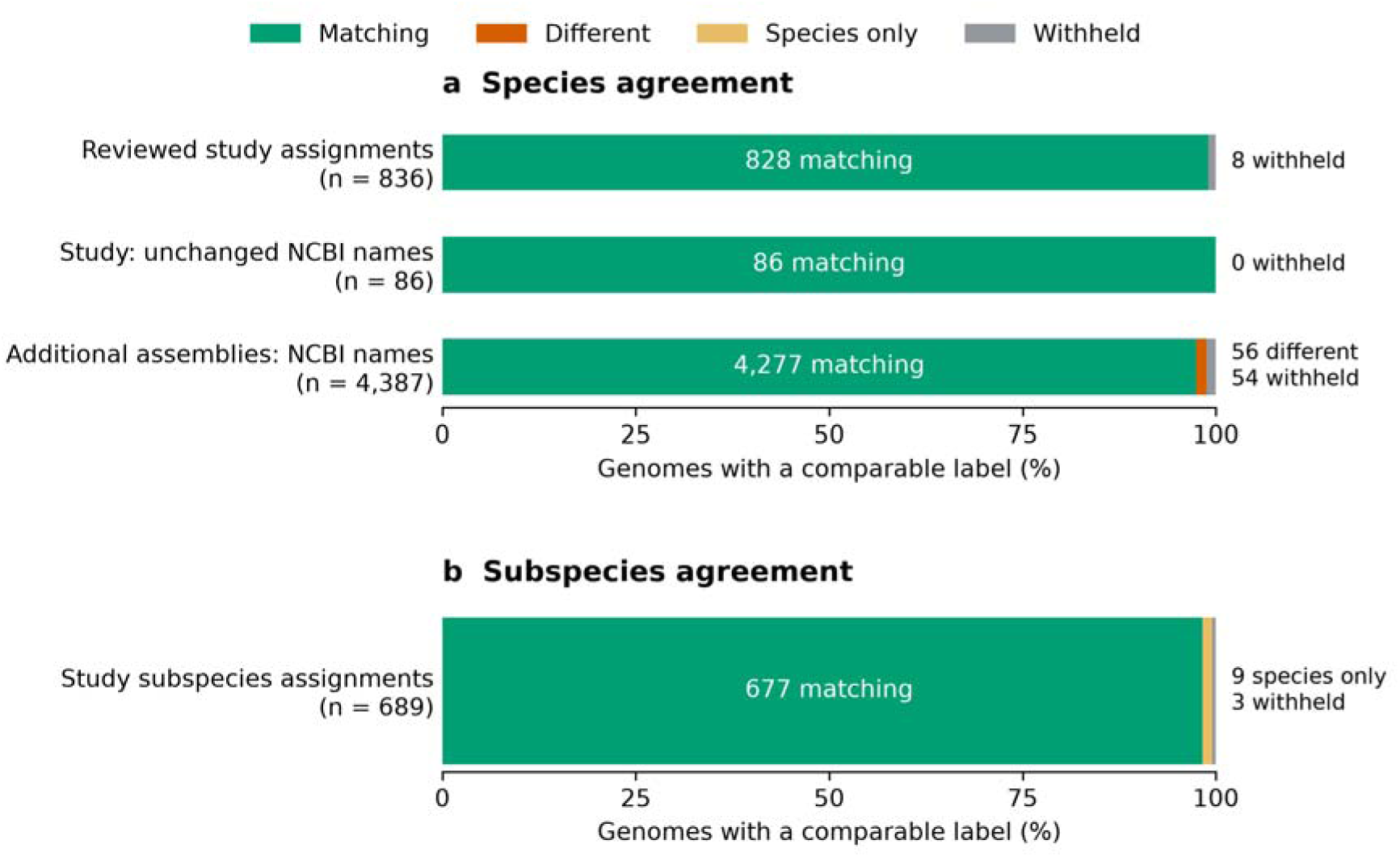
Agreement between Pterobacter species assignments and study classifications or NCBI names. (a) The first row compares 836 study genomes with assignments reviewed individually in this analysis. The second compares 86 study genomes with NCBI names that were compatible with the revised classification and were not individually reassessed. The third compares 4,387 additional public assemblies with their deposited NCBI species names. (b) Subspecies outcomes for the 689 study genomes with a subspecies expectation: 677 matching assignments, nine species-only assignments and three with no species assignment. Each bar includes every genome with a comparable label at that rank, including withheld outcomes. Numbers give genome counts; bar lengths give percentages within the stated denominator. The primary panel contained 140 references and queries were classified without tree placement. These comparisons measure agreement with study assignments or deposited names, rather than independent accuracy. Reference-removal experiments are shown separately in Figure S11.

**Table 3.** Agreement between Pterobacter and comparison labels.

| Evaluation set | Comparable genomes | Matching species assignments | Different species assignments | Withheld |
|---|---|---|---|---|
| Study: individually reviewed assignments | 836 | 828 | 0 | 8 |
| Study: unchanged NCBI names | 86 | 86 | 0 | 0 |
| Additional public assemblies | 4,387 | 4,277 | 56 | 54 |

The first study row uses assignments reviewed individually in this analysis. The second uses NCBI names compatible with the revised classification but not individually reassessed. The additional public assemblies are compared with their deposited NCBI names. The study rows together account for 922 named genomes among 946 non-reference study queries; the other 24 are unnamed or contextual genomes. The public comparison excludes 36 previously used development assemblies and 434 without a scorable species expectation from the 4,857 evaluated assemblies. These are comparisons with study assignments or deposited names, not independent accuracy estimates. Supplement S08 describes the outcome-selected reviews and reference-removal experiments. Counts describe the original classifier evaluation; the evaluated Rust implementation retained the native assignments.

We tested Pterobacter’s behaviour when given genomes for which there was no close reference by removing specific groups from the reference panel. In 123 comparisons with the expected subspecies removed, skani fell below the 96% species threshold but the reciprocal ANIm exceeded it with sufficient coverage, enabling a species assignment. Another 55 classification attempts remained unresolved because other reference comparisons showed inconsistent alignment coverage or conflicting results between reciprocal alignments, despite at least one qualifying ANIm match. (Figure S11; Supplement S08).

For genomovars, removing each query from its own reference group retained 148/173 source-group matches. Removing an entire group exposed a different problem: twelve previously ambiguous query comparisons then matched only a neighbouring group. We recorded these twelve neighbouring-group matches to demonstrate the effect of missing reference groups, but did not report them as genomovar assignments. Thus, an apparently unique match can depend on the breadth of diversity represented (Supplement S08; Figure S12).

Median elapsed time was 1.72 s for the skani-based route and 38.14 s across fifteen runs that included alignment-based ANI. Computational details are given in Supplement S10.

## Discussion

Our analysis revises the classification of a public *Enterobacter* collection and provides a way to apply the resulting species and subspecies assignments to new assemblies. The largest naming change concerns the lineages historically grouped within *E. hormaechei*. We retain the species containing its type strain and place the oharae, steigerwaltii, xiangfangensis and hoffmannii lineages within an expanded *E. xiangfangensis*. This classification retains the distinct lineages as subspecies within a broader species supported by type-strain comparisons, population-level ANI and phylogenetic analyses. We also propose reclassifying *E. quasiroggenkampii* and *E. rongchengensis* as subspecies of *E. roggenkampii* and *E. chengduensis*, respectively. The genomovars provide a finer description of diversity within the proposed *E. xiangfangensis* subsp. *hoffmannii*. The 99.5% threshold follows the within-species ANI gap reported by Rodriguez-R *et al.,* who describe genomovars as genomic groups that can contain multiple strains [31].

Our assessment here differs from both the broad *E. hormaechei* arrangement of Sutton *et al.* and the separation of *E. hoffmannii* and *E. xiangfangensis* supported by Wu *et al.* and De Maayer *et al.* [3, 5, 10]. Our expanded *E. xiangfangensis* is supported by concordant evidence from type-strain comparisons, population-level genomic similarity and the three phylogenetic analyses. All nine ANIm comparisons between the hoffmannii-associated types and the oharae, steigerwaltii and xiangfangensis types exceeded 95.9% identity with minimum aligned fractions of at least 84.8%. Across the wider collection, 98.2% of the 75,774 comparisons between these lineages met both the 95% ANI and bilateral 80% alignment-coverage criteria, showing that this relationship extends beyond the type strains. Retaining subspecies allows continued recognition of the differentiated groups within this broader species. The additional intestinihominis and nototheniae types shared 99.09–99.12% ANI with one another and the hoffmannii type. This high genomic similarity, together with the absence of a clear three-way population division corresponding to these names, supported their inclusion within a single subspecies. Together, these observations support the proposed classification while acknowledging that closely related lineages can admit alternative interpretations of species rank.

ANI and phylogenetics answer complementary parts of this problem. ANI quantifies sequence similarity between genomes and provides a basis for thresholds informed by observed gaps in genomic diversity. Large surveys support a species boundary near 95% ANI and a finer division within many species near 99.5% ANI [15, 31]. Phylogenetic analyses infer evolutionary relationships among genomes based on sequence variation. We compared phylogenies inferred from conserved markers, genome alignments and k-mer similarity to assess whether the proposed groups were consistently recovered across methods with different assumptions and sources of sequence information. Conflicting placements prompted further analyses of individual core genes, their genomic distribution and the influence of marker selection, revealing relationships that remained uncertain despite strong support in some individual trees. The unresolved *E. ludwigii* placement and alternative relationships among the hoffmannii-associated types limit conclusions about their evolutionary history, with none of the methods consistently favouring one phylogenetic approach over the others.

Some rank decisions required evidence beyond a single threshold. We retain *E. bugandensis* and *E. nematophilus* as separate species despite ANI overlap because similarity was higher within each group than between them, and the groups remained distinct across local trees. Additionally, *E. nematophilus* was initially represented only by its type genome; expanded sampling included six genomes, however this still left its diversity less well sampled than that of *E. bugandensis*. Conversely, the type ANIm and shared phylogenetic groups support the proposed subspecies within *E. roggenkampii* and *E. chengduensis*, despite published dDDH estimates below 70% [5, 33]. ANI and dDDH thresholds do not always yield equivalent species assignments, and published studies have demonstrated cases in which a fixed 70% dDDH boundary conflicts with broader evidence of genomic cohesion [34, 35]. We therefore interpreted these measures alongside population-level genomic comparisons and phylogenetic relationships. Our proposed subspecies treatments reflect this combined evidence, while the previously reported dDDH values remain as evidence supporting an alternative classification as separate species.

We developed Pterobacter so that users can apply these species and subspecies assignments directly to genome assemblies. Pterobacter uses skani to rapidly estimate ANI before conditionally running the slower but more accurate ANIm when skani is ambiguous (≥94.5% but <96%). This approach ensures speed in addition to accuracy for borderline cases, with a deliberately conservative threshold for species assignment to avoid incorrect calls. Across 5,803 study and public genomes, Pterobacter assigned a species to 5,671 (97.73%) and left 132 unassigned. Of the unassigned genomes, 62 had comparable named-species expectations (eight study genomes and 54 public genomes). This represents a highly feasible approach to reproducibly classify *Enterobacter* genomes in both research and clinical settings.

The Kraken2 results demonstrate how reference-database classifications can obscure distinctions between *Enterobacter* lineages. Kraken2 returned *E. hormaechei* subsp*. xiangfangensis* for sequences from several groups assigned to different subspecies in our analysis. Such assignments could confound comparisons of lineage distributions and associated traits across studies. Our classification provides an explicit genomic basis for distinguishing these groups, integrating type-strain relationships, population-level ANI and evidence from three phylogenetic approaches. Its advantage lies in the documented connection between names and genomic groups; the Kraken comparison demonstrates disagreement with this framework rather than an independent estimate of classification accuracy. The revised names, type-strain evidence and table linking each public genome to its original and revised assignments provide a basis for applying this classification consistently in future *Enterobacter* studies.

This study has several limitations. The taxonomic reassessment primarily used public assemblies deposited as complete genomes under *Enterobacter*, supplemented with selected type-strain genomes. This sampling does not capture the full diversity of the genus, particularly lineages represented mainly by draft assemblies or deposited under other genus names, and cannot establish the prevalence of taxa in clinical or environmental populations. Some proposed boundaries were supported by relatively few genomes. Broader sampling could reveal intermediate diversity and refine the species, subspecies and genomovar assignments.

Secondly, the genomic evidence did not support a single interpretation of every relationship or taxonomic boundary. The placement of *E. ludwigii* and the branching order of the hoffmannii-associated types remained unresolved, and the proposed quasiroggenkampii and rongchengensis subspecies treatments also conflict with published dDDH estimates, with additional alignment-coverage limitations for chengduensis–rongchengensis. No new dDDH measurements or phenotypic experiments were performed. As such, these treatments represent interpretations of the combined evidence rather than boundaries established uniformly by every measure.

In conclusion, our genomic reassessment supports revised species and subspecies boundaries within *Enterobacter*, including an expanded *E. xiangfangensis* that retains distinct lineages at subspecies rank. By integrating type-strain comparisons, population-level ANI and complementary phylogenetic analyses, we provide an explicit basis for these taxonomic proposals while identifying relationships that remain unresolved. Pterobacter makes this classification accessible for genome identification, with assignments withheld when reference evidence is insufficient. Together, the revised taxonomy, curated references and software provide a foundation for more consistent identification and comparison of *Enterobacter* lineages across studies.

### Proposed nomenclatural treatments

We propose five transferred combinations and three corresponding type-subspecies combinations (Table 4), together with emended descriptions of *E. xiangfangensis* and *E. hormaechei*. Phenotypic properties are retained from the cited original descriptions. Genome sizes and G+C ranges describe the assemblies examined here and are not diagnostic limits for membership. Supplement S11 records type-strain documentation and the limits of the competing-name screen.

**Table 4.** Proposed taxonomic combinations and type genomes.

| Proposed name (E. prefix) | Act | Designated type collections | Whole type genome | bp; G+C mol% | Assigned genomes |
|---|---|---|---|---|---|
| xiangfangensis subsp. *oharae* | comb. nov. | CIP 108490; DSM 16687; EN-314 | GCA_001729705.1 | 4,724,316; 55.58 | 18 |
| xiangfangensis subsp. *steigerwaltii* | comb. nov. | CIP 108489; DSM 16691; EN-562 | GCA_001729725.1 | 4,782,480; 55.55 | 256 |
| xiangfangensis subsp. *hoffmannii* | comb. nov. | EN-114; DSM 14563; CCM 9140 (LMG 30171 documentation not relied on) | GCA_001729745.1 | 4,678,566; 55.33 | 173 |
| roggenkampii subsp. *quasiroggenkampii* | comb. nov. | GDMCC 1.1742; KCTC 52992; WCHECL1060 | GCA_001039365.2 | 4,807,091; 55.69 | 6 |
| chengduensis subsp. *rongchengensis* | comb. nov. | 170250; GDMCC 1.3670; JCM 36189 | GCA_034008275.1 | 4,952,144; 56.18 | 2 |
| xiangfangensis subsp. *xiangfangensis* | type subspecies; comb. nov. | 10-17; CCUG 62994; LMG 27195; NCIMB 14836 | GCA_001729785.1 | 4,661,849; 55.28 | 163 |
| roggenkampii subsp. *roggenkampii* | type subspecies; comb. nov. | DSM 16690; EN-117; LMG 30172 | GCA_001729805.1 | 4,899,997; 56.04 | 81 |
| chengduensis subsp. *chengduensis* | type subspecies; comb. nov. | CCTCC AB 2017105; CCUG 76941; GDMCC 1.1216; KCTC 52993; WCHECl-C4 | GCA_001984825.2 | 5,218,125; 55.74 | 3 |

The table contains five proposed transfers and three corresponding type-subspecies combinations. Counts include assigned study genomes and any reference-only types; the peripheral xiangfangensis species-only member is outside the subspecies counts. Type-strain culture-collection identifiers and supporting nomenclatural records are provided in Table S21. Genomovars have no nomenclatural standing.

*Enterobacter xiangfangensis* subsp. *oharae* (Hoffmann et al. 2016) comb. nov.

Etymology: o’ha.rae. N.L. gen. n. oharae, honoring Caroline M. O’Hara, an American microbiologist. Basonym and homotypic synonym: *Enterobacter hormaechei* subsp. *oharae* Hoffmann et al. 2016. No heterotypic synonym is proposed. The properties remain those described by Hoffmann et al. [7, 36]. The oharae lineage is retained as a subspecies within the expanded *E. xiangfangensis*.

The type strain is EN-314 (=DSM 16687 =CIP 108490), originally obtained from a mouth swab of a two-year-old infant. Its complete type genome is GCA_001729705.1, comprising 4,724,316 bp with G+C 55.58 mol%. The 18 assigned genomes range from 4.72 to 5.22 Mb and 54.87–55.60 mol% G+C.

*Enterobacter xiangfangensis* subsp. *steigerwaltii* (Hoffmann et al. 2016) comb. nov.

Etymology: stei.ger.wal’ti.i. N.L. gen. masc. n. steigerwaltii, honoring Arnold G. Steigerwalt, an American microbiologist. Basonym and homotypic synonym: *Enterobacter hormaechei* subsp. *steigerwaltii* Hoffmann et al. 2016. No heterotypic synonym is proposed. The description refers to the original properties [7, 36], with transfer into the expanded *E. xiangfangensis* and retention of the steigerwaltii type-associated subgroup.

The type strain is EN-562 (=DSM 16691 =CIP 108489), isolated from an infected surgical skin wound. Type genome GCA_001729725.1 comprises 4,782,480 bp with G+C 55.55 mol%. The 256 assigned genomes range from 4.55 to 5.67 Mb and 54.41–55.77 mol% G+C.

*Enterobacter xiangfangensis* subsp. *hoffmannii* (Sutton et al. 2018) comb. nov.

Etymology: hoff.man’ni.i. N.L. gen. masc. n. hoffmannii, honoring Harald Hoffmann, a German microbiologist who studied the phylogenetic structure of the *E. cloacae* complex. Basonym: *Enterobacter hormaechei* subsp. *hoffmannii* Sutton et al. 2018 [10, 37].

The homotypic synonyms are *E. hormaechei* subsp. *hoffmannii* Sutton et al. 2018; *E. hoffmannii* (Sutton et al. 2018) Sedláček et al. 2026; *E. hoffmannii* subsp. *hoffmannii* (Sutton et al. 2018) Sedláček et al. 2026; and the effectively published, non-valid “*E. hoffmannii*” (Sutton et al. 2018) Wu et al. 2020 [5, 12, 14]. Proposed heterotypic synonyms are *E. intestinihominis* Hitch et al. 2025 and *E. hoffmannii* subsp. *nototheniae* Sedláček et al. 2026 [11, 13, 14].

The description follows Sutton et al., with circumscription extended to the intestinihominis type CLA-AC-H004 (GCA_040096145.1) and nototheniae type P5473 (GCA_052922205.1). These genomes form one subspecies within *E. xiangfangensis*, with uncertainty retained about the branching history of the three types. Its 25 genomovars are informal subdivisions without nomenclatural standing.

The type strain is EN-114 (=DSM 14563 =CCM 9140), isolated from a patient’s respiratory tract. Type genome GCA_001729745.1 comprises 4,678,566 bp with G+C 55.33 mol%. The 173 assigned study/reference genomes range from 4.54 to 5.56 Mb and 54.29–55.42 mol% G+C. VL230 did not receive documentation for the additional LMG 30171 deposit; that number is not relied on here.

*Enterobacter roggenkampii* subsp. *quasiroggenkampii* (Wu et al. 2020) comb. nov.

Etymology: qua.si.rog.gen.kam’pi.i. Latin quasi, nearly or almost; quasiroggenkampii, almost roggenkampii. Basonym and homotypic synonym: *Enterobacter quasiroggenkampii* Wu et al. 2020. No heterotypic synonym is proposed. The original description is retained by reference [5, 38]. Its type-associated lineage is placed at subspecies rank within the expanded *E. roggenkampii* species. The published 64.8% dDDH and 95.37% ANI to the roggenkampii type remain part of the original evidence; the present type ANIm, population comparisons and tree context support the stated alternative rank opinion without claiming universal agreement.

The type strain is WCHECL1060 (=GDMCC 1.1742 =KCTC 52992), recovered from a blood culture in Chengdu, China. Draft genome GCA_001039365.2 comprises 4,807,091 bp with G+C 55.69 mol%. The six assigned genomes range from 4.70 to 5.47 Mb and 55.28–56.00 mol% G+C.

*Enterobacter chengduensis* subsp. *rongchengensis* (He et al. 2025) comb. nov.

Etymology: rong.cheng.en’sis. N.L. masc. adj. rongchengensis, pertaining to Rongcheng, another name for Chengdu, China. Basonym and homotypic synonym: *Enterobacter rongchengensis* He et al. 2025. The effective description was published in 2024 and validated in 2025 [33, 39]. No heterotypic synonym is proposed.

The properties are those of the original description, with the two-member lineage treated as a subspecies within chengduensis. The original Table 1 reports 95.58% ANI and 64.4% dDDH against the chengduensis type. Its prose transposes the strain comparison; the table is the numerical source. The merger retains the observed chromosome-scope AF failures and does not claim that the published dDDH supports the same rank.

The type strain is 170250 (=GDMCC 1.3670 =JCM 36189), recovered from human blood in Chengdu, China. Draft genome GCA_034008275.1 comprises 4,952,144 bp with G+C 56.18 mol%. The two assigned genomes range from 4.95 to 4.97 Mb and 56.10–56.18 mol% G+C.

Description of the type subspecies *Enterobacter xiangfangensis* subsp. *xiangfangensis* (Gu et al. 2014) comb. nov.

Etymology: xi.ang.fang.en’sis, pertaining to Xiangfang, a district of Harbin, Heilongjiang Province, China. Basonym: *E. xiangfangensis* Gu et al. 2014. Homotypic synonym: *E. hormaechei* subsp. *xiangfangensis* (Gu et al. 2014) Sutton et al. 2018 [9, 10, 37].

The original species description is retained, restricted to the subgroup containing its type. The type strain is 10-17 (=CCUG 62994 =LMG 27195 =NCIMB 14836), isolated from Chinese traditional sourdough. Genome GCA_001729785.1 comprises 4,661,849 bp with G+C 55.28 mol%. The 163 assigned genomes range from 4.51 to 5.57 Mb and 54.33–55.58 mol% G+C.

Description of the type subspecies *Enterobacter roggenkampii* subsp. *roggenkampii* (Sutton et al. 2018) comb. nov.

The etymology and properties refer to the species description [10, 37], restricted to the type-containing subgroup. Basonym and homotypic synonym: *E. roggenkampii* Sutton et al. 2018. The type strain is EN-117 (=DSM 16690 =LMG 30172), isolated from human stool. Genome GCA_001729805.1 comprises 4,899,997 bp with G+C 56.04 mol%. The 81 assigned genomes range from 4.63 to 5.69 Mb and 55.23–56.16 mol% G+C.

Description of the type subspecies *Enterobacter chengduensis* subsp. *chengduensis* (Wu et al. 2019) comb. nov.

The etymology and properties refer to the species description [40], restricted to the type-containing subgroup. Basonym and homotypic synonym: *E. chengduensis* Wu et al. 2019. The type strain is WCHECl-C4 (=CCTCC AB 2017105 =CCUG 76941 =GDMCC 1.1216 =KCTC 52993), isolated from human blood in Chengdu, China. Genome GCA_001984825.2 comprises 5,218,125 bp with G+C 55.74 mol%. The three assigned genomes range from 5.09 to 5.61 Mb and 55.18–56.14 mol% G+C.

### Emended description of *Enterobacter xiangfangensis* Gu et al. 2014

The original properties [9] are retained with a broadened circumscription comprising the proposed type subspecies and oharae, steigerwaltii and hoffmannii. This includes genomes previously placed under the corresponding hormaechei subdivisions, *E. intestinihominis* and *E. hoffmannii* subsp. *nototheniae*. The collection contains 611 assigned study/reference genomes, including one peripheral member assigned only to species; whole-assembly size is 4.51–5.67 Mb and G+C 54.29–55.77 mol%. The type remains 10-17; type genome GCA_001729785.1.

### Emended description of *Enterobacter hormaechei* O’Hara et al. 1990

The original properties [8] are retained for the lineage containing the *E. hormaechei* type. The oharae, steigerwaltii, xiangfangensis and hoffmannii subdivisions are transferred into the adopted xiangfangensis circumscription. Subsp. hormaechei remains the retained type subspecies; the binomial is used here. The thirteen assigned genomes range from 4.55 to 5.37 Mb and 54.64–55.46 mol% G+C. The type is 0992-77 (=ATCC 49162 =DSM 12409); genome GCA_019048245.1 comprises 4,915,930 bp with G+C 55.14 mol%.

### Other nomenclatural opinions and unresolved names

Quasimori is proposed as a later heterotypic synonym of mori, and dykesii as a later heterotypic synonym of asburiae; these are taxonomic opinions without creation of a new name. Existing cloacae subspecies names are retained. “*E. adelaidei*” remains quoted and not validly published; no validation attempt is proposed. Unnamed lineages and informal genomovars receive no formal new names. The four names without selected type assemblies and the limits of the siamensis priority comparison are detailed in Supplement S11 [30, 41].

## Data and code availability

The public assembly accessions, original labels and revised assignments are provided in Table S01. Source data and scripts reproduce the main figures and tables. The software evaluation used source commit 834f4c288e9c163f316f3668f0219004d0adc6da, before adoption of the Pterobacter name; the executable reported terobacter 0.0.0. Supplement S10 records the evaluated software and reference versions separately from subsequent releases. The reference database (version 2026-10-15) is available at https://doi.org/10.6084/m9.figshare.34640883.v1 [42]. Analysis scripts, environments and source data are available at https://github.com/mbhall88/Enterobacter-analysis (commit 3a2baebaa1c82149451fe6039659bfc83ec1ca5a); the analysed sequences, alignments, trees and frozen evaluation packages are archived at https://doi.org/10.6084/m9.figshare.34742013.v1. The revised supplementary tables and indexed workbook accompany this manuscript; the earlier table version is archived at https://doi.org/10.6084/m9.figshare.34742010.v1. Pterobacter source and release 0.1.0 are available at https://github.com/mbhall88/pterobacter/releases/tag/0.1.0. The later public release is distinct from the version used for the biological evaluation.

## Author statements

Author contributions:

Monique Clifton: [Data curation; Formal analysis; Investigation; Methodology; Visualization; Writing – original draft; Writing – review & editing].

Jocelyn Hume: [Investigation; Methodology; Resources; Supervision; Writing – review & editing].

Michael B Hall: Conceptualization; Data curation; Formal analysis; Investigation; Methodology; Software; Validation; Visualization; Writing – original draft; Writing – review & editing.

Leah W Roberts: [Conceptualization; Funding acquisition; Investigation; Methodology; Project administration; Resources; Supervision; Visualization; Writing – review & editing].

Funding: LWR is supported by a National Health and Medical Research Council (NHMRC) Investigator grant [Grant ID: 2026911].

Conflicts of interest: The authors declare that there are no conflicts of interest.

Ethical approval and consent: This study reanalysed publicly available genome assemblies and involved no human participants or animal experiments; ethical approval and consent to participate were not required.

## Acknowledgements

We would like to acknowledge Jake Lacey for engaging in initial discussion regarding the project. This work was supported by resources provided by the University of Queensland Research Computing Centre’s Bunya supercomputer [43], with funding from the University of Queensland, Brisbane, Australia.

## Notes

https://doi.org/10.6084/m9.figshare.34640883.v1

https://doi.org/10.6084/m9.figshare.34742013.v1

https://doi.org/10.6084/m9.figshare.34742010.v1

https://github.com/mbhall88/pterobacter/releases/tag/0.1.0

https://github.com/mbhall88/Enterobacter-analysis

### Competing Interest Statement

The authors have declared no competing interest.

