## Supplementary Materials for "Genomic reassessment of *Enterobacter* species and subspecies"

### Supplementary material

This supplement describes genome selection and sequence preparation (S01), ANI and type-strain comparisons (S02), phylogenetic analyses (S03–S04), sequence-selection sensitivities (S05), taxonomic interpretations (S06), genomovar definitions (S07), classifier evaluation (S08), the Kraken comparison (S09), software and computational measurements (S10), nomenclatural evidence (S11), and public database and supporting files (S12). References are listed in the main bibliography; numbered citations use the same reference numbers in both documents. Supplementary tables provide accession-level data and the complete comparison results.

#### S01. Genome selection and sequence preparation

##### Study collection and accession catalogue

The initial collection contained 1,079 NCBI assemblies submitted as *Enterobacter* and designated complete on 5 February 2026. Paired GenBank and RefSeq accessions were recorded as representations of the same assembly rather than counted twice. Identification of duplicate submissions used accession and provenance information; identical sequence or a shared strain label alone did not establish isolate identity. The collection was restricted to records submitted under *Enterobacter* and therefore did not comprehensively sample draft genomes or organisms submitted under other genus names.

Four records were identified as duplicate submissions of other included assemblies, and one assembly was excluded because its quality and taxonomic reliability were inadequate. These include GCA_023547225.1, GCA_023547315.1, GCA_052982295.1 and GCA_052982515.1, represented by retained assemblies GCA_051131905.1, GCA_051131895.1, GCA_030411175.2 and GCA_030411295.2, respectively. GCA_042434945.1 was excluded because NCBI flagged contamination and failed annotation completeness, marker recovery was poor and its taxonomic placement was inconclusive. Its CheckM completeness estimate was 92.65%. These accession-specific decisions are recorded in Table S01. This left 1,074 study assemblies.

Four assemblies were retained in the study catalogue but excluded from the final comparison of classifications. GCA_002356475.1 had frameshift-associated marker failure; GCA_041890495.1 was chimeric; GCA_006385655.1 had 12.81% estimated completeness; and GCA_050242595.1 was excluded following a further quality assessment (61 missing marker families). Their available ANI measurements were retained, but no final naming comparison was made for these four. The remaining 1,070 assemblies form the denominator for Table 1 and the Kraken comparison.

Table S01 contains 1,098 accession records, with the following composition. The five initial exclusions are retained in the table so that the selection can be reconstructed; they are outside the 1,074-genome study set. The diagnostic addition was used in a focused comparison and was not added to the study denominator.

| Record category | Accessions | Relationship to the study collection |
| --- | --- | --- |
| Study assemblies | 1,074 | Includes the four later sequence-quality exclusions; also includes 13 out-of-genus context genomes retained in the principal 1,089-tip comparison but removed for the 1,076-tip ingroup analysis |
| Additional type-strain genomes | 16 | Absent from the study collection |
| Outgroups | 2 | *Cronobacter sakazakii* and *Klebsiella aerogenes* |
| Initial exclusions | 5 | Four duplicate submissions and one unsuitable assembly |
| Diagnostic addition | 1 | BS0013, GCA_030510045.1 |
| Total catalogue | 1,098 | Accession records, not a single analysis cohort |

##### Chromosome and type-strain sequence inputs

The taxonomic analyses used chromosome sequences from complete genomes where possible. Contigs shorter than 1 Mb were removed using a python script and genomes checked for a single retained contig. The 33 selected type-strain assemblies were retrieved with NCBI Datasets 18.25.1 on 17 September 2026, including sequence reports. Seventeen were complete genomes (and thus not added as they were already in the study collection) and sixteen were drafts (thirteen at contig level and three at scaffold level). Draft types retained all contigs because their chromosome or plasmid origin could not be assigned reliably. Thus, a chromosome-labelled input for a draft does not imply a closed chromosome.

##### Analysis-specific subsets and recorded names

The first phylogenetic dataset omitted the frameshift-affected and chimeric study assemblies, giving 1,072 study genomes plus sixteen additional types and two outgroups (1,090 inputs). The very low-completeness assembly was subsequently pruned, giving the principal 1,089-tip comparison. Removing thirteen additional genomes found to be outside *Enterobacter* gave the 1,076-tip ingroup analysis. GCA_050242595.1 (which lacked 61 BAC120 marker families) was excluded later; the effect was examined by pruning the existing trees to 1,088 tips. These are successive analysis subsets, rather than conflicting estimates of the study collection size. Figure 1 relates them to the final 1,070-genome classification set and classifier evaluation.

For each accession, Table S01 records the original NCBI name, the name retrieved on 23 September 2026, the classification proposed here and any boundary or sequence-quality qualification. Only one of the 1,070 eligible assemblies changed its recorded NCBI name between the snapshots (February to September 2026): GCA_051302795.1 changed from *E. ludwigii* to Pantoea sp. E2023-21.

Table S01 reports the numbers of BAC120 marker families detected as single copies, missing, or present in multiple copies for each genome in the initial 1,090-genome phylogenetic dataset. Genomes missing five or more marker families are flagged to aid interpretation of their phylogenetic placement. This flag did not automatically trigger exclusion, alter taxonomic assignments or determine eligibility for Pterobacter classification. Genomes not included in this marker analysis are marked as unassessed.

#### S02. ANI screening and type-strain comparisons

The initial screen compared every study chromosome set with every selected type-strain sequence set. Skani 0.3.1 was run using dist --min-af 0 -s 0 --detailed, with query and reference lists supplied through --ql and --rl, and four threads per call. Setting the output filters to zero retained comparisons below the later analytical thresholds. All 35,442 study–type pairs were measured in each orientation. Candidate species relationships required ANI ≥95% and AF ≥80% for both genomes. AF measures the fraction of each input genome represented in the comparison. Requiring coverage of both inputs avoids qualifying a pair on only a small shared region.

Selected boundary comparisons were repeated with pyANI [16] using NUCmer 3.1/MUMmer 3.23 [44]. The lower ANI of the two orientations was retained, and coverage was measured from merged alignment intervals for both genomes in each orientation. This coverage differs from skani’s estimate based on chained matches. BLAST-based ANI was additionally measured for the *E. cloacae*–*E. dissolvens* types. Published dDDH values were used as external context, without new dDDH computation.

Table S03 links the Figure 2 identifiers to the 33 type accessions and two outgroups, their historical type names and the proposed classifications. Alternative assemblies carrying equivalent type-strain identifiers were not treated as interchangeable unless their origin and identity could be verified. For *E. kobei*, the selected DSM 13645 genome was retained; a divergent JCM-labelled assembly has unresolved culture history. Two *E. sichuanensis* assemblies carrying the WCHECL1597 label share about 98.18% ANI, so their common culture label does not establish that they represent the same isolate. Their alternative placements were examined without silently substituting one assembly for the other.

#### S03. Comparison of the three phylogenetic methods

Sequences were prepared for Verticall 0.4.2 by removing non-ACGT bases and splitting records at those positions. The same prepared sequences were used for all three phylogenetic approaches. For the conserved-marker tree, GTDB-Tk 2.7.2 with reference release R232 identified and aligned the BAC120 marker proteins. Alignment positions containing standard amino acids in at least 80% of genomes were retained, and FastTree 2.2.0 inferred the tree using the WAG substitution model with gamma rescaling. For the genome-alignment tree, Verticall 0.4.2 estimated distances from vertically inherited sequence, and FastME 2.1.6.3 inferred a balanced minimum-evolution tree using nearest-neighbour interchange and subtree pruning and regrafting. For the k-mer approach, Mashtree 1.4.6 used Mash 2.3 and quicktree 2.5 with a k-mer length of 21 and 10,000 hashes per genome. We summarised 1,000 sketch-seed replicate trees as a majority-rule consensus, with branch lengths averaged across replicates containing each branch.

Before comparing trees, we collapsed branches at or below the method-specific length thresholds: 2.5180037 × 10⁻⁵ substitutions per site for BAC120 (1/39,714 alignment columns), 2.0915 × 10⁻⁷ for Verticall/FastME (one base across the median 4,781,248-bp chromosome), and 2.381 × 10⁻⁶ for Mashtree (one differing hash at k = 21 and 10,000 hashes). Retained BAC120 branches also required SH-like local support ≥0.95. FastME had no support filter; the Mashtree majority-rule consensus already retained only splits present in more than half of the seed replicates. These thresholds are not equivalent biological distances. Each retained split was classified as shared, unresolved or contradicted in the other tree; backbone splits contained at least ten genomes on each side. Table S04 reports the comparisons using these method-specific rules and length-only filters. The latter still used the majority-rule Mashtree consensus.

Sensitivity comparisons examined alternative marker models and implementations, including double precision; alternative Verticall distances; and Mashtree seed choice, replicate count and sketch size. We also compared the primary occupancy-filtered marker alignment with the canonical GTDB mask and assessed genome exclusions by pruning existing trees. These checks examine different sources of variation and do not provide a common scale of phylogenetic accuracy (Table S04).

All methods used the same prepared sequences. Removing 7,554 non-ACGT bases affected seven genomes and created 128 additional sequence fragments. Subsequent genome exclusions were assessed by removing tips from existing trees without recalculating alignments, branch lengths or support; the Mashtree consensus was checked separately.

For Figure 2, each 35-tip projection was rooted halfway along the existing edge separating Cronobacter sakazakii and Klebsiella aerogenes from the 33 type-strain tips. This changes only the display orientation, preserving unrooted splits, pairwise path lengths and support values. The two outgroups could not jointly root the full 1,089-tip trees; joint rooting was possible in the 1,076-tip scope after excluding thirteen out-of-genus context genomes. The type-strain display therefore does not resolve the genus placement of those additional genomes.

#### S04. Focused investigation of conflicting tree relationships

We constructed three focused datasets to examine the placement of E. ludwigii relative to E. cloacae, E. asburiae, E. roggenkampii and E. kobei; the placement of E. sichuanensis relative to E. chinensis and E. cloacae; and the branching order of the E. hoffmannii, E. intestinihominis and E. hoffmannii subsp. nototheniae type strains. Pyrodigal 3.7.1 and Panaroo 1.8.0 identified core genes, and MAFFT 7.526 aligned them [25, 26, 27]. The focused datasets (above) contained 3,209–3,787 codon-aligned genes. We compared gene trees (made using IQ-TREE 3.1.1 [28]), gene and site concordance, quartet patterns and the genomic positions of supporting genes [45]. Each quartet comprised four selected genomes, for which there are three possible unrooted branching arrangements. For each comparison, we counted the gene trees supporting each arrangement and expressed these counts as proportions of trees containing all four genomes and resolving their relationship. Trees lacking a selected genome or leaving the quartet unresolved were excluded from that comparison’s denominator. Approximately unbiased (AU) tests of constrained topologies used 10,000 RELL replicates [46]. The following sections distinguish the hypotheses actually tested from the unresolved or failed analyses.

##### E. ludwigii

The closest relatives of E. ludwigii remained unresolved. The BAC120 marker tree placed it with E. cloacae, whereas the distance-based trees associated it with the group containing E. asburiae, E. roggenkampii and E. kobei. Individual core-gene trees supported both placements, showing that the disagreement extended beyond the BAC120 markers.

We assessed these competing relationships using four-genome comparisons, or quartets. Across comparisons using E. asburiae or E. kobei as representatives of the alternative lineage, the two placements together accounted for 71–84% of resolved gene-tree quartets. Comparisons using E. sichuanensis as the alternative representative gave combined proportions of 66–75%. These percentages describe support for the two placements combined; neither placement was consistently favoured across gene sets and representatives. Genes supporting the same placement often occurred next to one another along the genome, indicating that support was concentrated in genomic regions rather than distributed among independent genes.

Some topology tests could not be used to resolve this conflict. In the expanded genome panel, the constraint intended to reproduce the BAC120 placement instead forced E. ludwigii to branch outside all other sampled Enterobacter lineages. This differed from its actual position within the 402-genome clade containing E. ludwigii and E. cloacae. Results from this incorrect constraint, including its use in the expanded marker analyses, were therefore excluded as evidence against the BAC120 placement. In the original, smaller panel, the correctly specified marker placement was the highest-likelihood tree. A corrected test on the expanded panel was not completed (Table S05).

##### E. sichuanensis and E. chinensis

The broader core-gene evidence favoured a relationship between E. sichuanensis and E. chinensis. Verticall/FastME and Mashtree recovered this pairing, whereas the BAC120 tree placed E. sichuanensis with E. cloacae (Figures S4 and S6). Across the tested four-genome comparisons, 65.7–78.8% of resolved core-gene quartets supported the sichuanensis–chinensis pairing. A topology test also rejected the alternative sichuanensis–cloacae pairing under the tested conditions (conditional p-AU <0.001).

The genomic distribution of the BAC120 markers offered a possible explanation for their different result. These markers were disproportionately located in regions supporting the sichuanensis–cloacae relationship (descriptive Fisher test, p = 1.8 × 10⁻⁵). However, neighbouring genes may share the same evolutionary history, so treating them as independent observations could overstate the statistical evidence. We did not perform a block-permutation test to account for this dependence.

We also examined an alternative assembly, GCA_025002605.1, carrying the same WCHECL1597 strain label as the selected E. sichuanensis type assembly, GCA_002939185.1. The two assemblies shared approximately 98.18% ANI and had similar broad phylogenetic placements. These observations did not establish that they represented the same isolate, so the alternative assembly did not replace the selected type genome.

##### E. hoffmannii, E. intestinihominis and the nototheniae type

The branching order of the three type strains remained unresolved. We compared three possibilities: hoffmannii–intestinihominis, hoffmannii–nototheniae, and intestinihominis–nototheniae, with the third type branching separately in each case. These analyses examined relationships within the proposed E. xiangfangensis subsp. hoffmannii; they did not test whether the three types should belong to different subspecies.

Individual core-gene trees provided almost equal support for the three pairings, with quartet proportions of approximately 34%, 32% and 34%. Site-based comparisons were also inconclusive. In the unmasked alignment, intestinihominis–nototheniae had the largest site-support share with each of three outgroups, but the ranking of the other pairings changed. Site-concordance values calculated using alternative reference trees did not consistently favour the same pairing.

Likelihood comparisons depended on which codon positions were included. Using all three positions, intestinihominis–nototheniae ranked first and both alternative pairings were rejected (conditional p-AU <0.001). Using only the first and second positions, hoffmannii–nototheniae was rejected (p-AU = 0.0119), but hoffmannii–intestinihominis was not (p-AU = 0.199). A second tested tree containing the intestinihominis–nototheniae pairing was also not rejected (p-AU = 0.20; Table S05).

These likelihood results require caution. Model parameters were held fixed from selected searches that had not converged, so the tests were conditional on those parameters and the candidate trees examined. Three expanded searches using all codon positions failed, and one planned AU test was not run. In addition, E. hormaechei was absent from 583 of 3,731 core-gene trees; comparisons requiring that genome therefore used only the available, resolved trees. Taken together, the conflicting signals and incomplete analyses prevented a confident choice among the three branching arrangements.

#### S05. Sensitivity to sequence selection and assembly structure

##### Records omitted from chromosome inputs

We examined 2,839 sequences (plasmids etc.) omitted when preparing chromosome-only inputs from the study assemblies (251.2 Mb in total) to determine whether they contained chromosome-like sequence that could affect the taxonomic comparisons. Records were aligned with NUCmer 3.1 (--maxmatch) to deposited chromosome and plasmid records of eligible neighbouring complete assemblies. After filtering at ≥95% identity, we measured the union of query intervals aligned to each target class independently. A record was chromosome-like if at least 50% aligned to chromosomes and less than 20% to plasmids; the reverse thresholds defined plasmid-like records. At least 20% to both classes defined shared or mixed matches; all other records were unresolved. These are sequence-similarity categories, not proof of physical location. This comparison screened 1,367 records from 748 assemblies: twenty records in nineteen assemblies were chromosome-like, 642 were plasmid-like, fifteen had shared or mixed matches, and 690 remained unresolved. The 1,472 unscreened records comprised 1,308 shorter than 50 kb and 164 without an eligible neighbour. Table S06 lists the records, comparison scope and classifications. An earlier marker search covered only 4.4% of excluded bases and was not treated as a complete screen.

Neighbours were chosen using GTDB species context, independently of the taxonomic treatments proposed here. Consequently, some comparisons that originally crossed differently named groups now fall within a proposed merged species. A chromosome-like match does not establish physical linkage or an assembly error. In four cases, excluded records matched the retained chromosome almost completely at approximately 99.9–100% identity (one unrounded value was 99.89945%). These records therefore largely duplicated retained sequence, rather than supplying chromosome regions missing from the analysis.

All sixteen draft type assemblies, comprising 891 contigs, were inventoried (Table S07). Fourteen had an eligible complete neighbour; *E. nematophilus* and *E. chinensis* did not. Fifteen candidate plasmid-like contigs occurred in *E. soli*, *E. chuandaensis* and *E. intestinihominis*. For nine genome pairs, we repeated ANIm comparisons using draft type-strain assemblies with and without candidate plasmid-like contigs. Three pairs met the ANI ≥95% and bilateral alignment-coverage ≥80% criteria before and after removal, while six remained below 95% ANI. All nine met the coverage criteria with both input versions. This result applies to the tested pairs and does not establish that plasmid removal would leave every comparison or the full phylogenies unchanged (Table S08).

##### Additional sequence in isolates matching the chromosome

Three records annotated as plasmids contained chromosome-associated sequence. One was the *E. ludwigii* record CP017280.1; adding it back to the analysis did not change the full or masked BAC120 alignment rows for this genome. The second record was an oharae RHBSTW-00844 (GCA_013731155.1). Its excluded record CP056156.1 was 361,638 bp long; 98.892% aligned to a neighbouring chromosome, and it contained eleven BAC120 families missing from the selected chromosome input. Adding CP056156.1 restored those families and added a duplicate, increasing uniquely recovered families from 108 to 118. The deposited plasmid annotation and sequence similarity do not establish its physical connection to the chromosome. With 51 tips and the same primary columns, LG+F+G4 model and support procedure, adding the omitted sequence preserved the 18-genome subgroup and its closest neighbouring group. However, 16 branching relationships changed, or 14 when only branches of positive length were counted (Table S09). The shorter GTDB-filtered alignment produced a poorly resolved tree in which the oharae group was not recovered as a distinct group.

The third record from an *E. hormaechei* CP077309.1 contained dnaA-dnaN-recF-gyrB and eleven BAC120 families absent from CP077308.1. Whole-assembly and selected-chromosome inputs yielded 117 and 108 uniquely recovered marker families, respectively; this net difference includes multicopy effects.

Replacing the selected type chromosome with the whole type assembly changed three outcomes. Among the 438 genomes associated with oharae, steigerwaltii and xiangfangensis, seven then passed the skani ANI/AF screen, compared with none using the selected chromosome; median query AF increased from 73.85 to 83.91%. The whole-assembly input includes every deposited record and redundant repeat ends, so this is not an isolated test of CP077309.1. Whole-type population ANIm was not performed. The alternative input was used for sensitivity analysis and did not replace the reference underlying the reported primary comparisons. The assembly was deposited as complete, but chromosome-associated sequence occurs in both CP077308.1 and CP077309.1. The sequence evidence therefore exposes a limitation of treating the selected record alone as the complete chromosome; it does not establish the physical topology of the other record.

#### S06. Evidence for individual taxonomic treatments

Figures S1, S4–S9 show the relevant type strains in each of the three phylogenies alongside type ANI and alignment coverage. Figure S2 shows the corresponding population comparisons for *E. xiangfangensis* and *E. hormaechei*. The plots retain the original sequence scopes and distinguish the historical type names from the proposed classifications.

##### *E. xiangfangensis* and *E. hormaechei*

The type-strain comparison is shown in Figure S1.

The expanded *E. xiangfangensis* comprises a 173-genome lineage containing the hoffmannii, intestinihominis and nototheniae types, and 438 genomes associated with the oharae, steigerwaltii and xiangfangensis types. The 173-genome lineage includes one additional type genome absent from the study collection. Including the thirteen-genome *E. hormaechei* type lineage gives the broader 624-genome clade. The *E. hormaechei* lineage is retained as a separate species, while the two larger sets are treated within *E. xiangfangensis* (main Table 2).

Of the 438 genomes, 18 were assigned to subsp. *oharae*, 256 to subsp. *steigerwaltii* and 163 to subsp. *xiangfangensis*. The peripheral GCA_039734725.2 received a species assignment only. Within the 437 subspecies-assigned genomes, ANI ranges were 98.18–100% within subspecies and 96.77–98.11% between subspecies. A separate sensitivity analysis assigned all 438 genomes to their nearest type for comparison, placing the peripheral genome in a descriptive steigerwaltii bin. The within-bin and between-bin minima were then 96.19 and 95.70%. That bin contained 257 genomes and 32,896 within-bin pairs, compared with 256 genomes, 32,640 pairs and minimum ANI 98.27% in the assigned steigerwaltii subspecies. Nearest-type bins are therefore not interchangeable with subspecies assignments. All three trees recovered the broader groups, although the smallest clade containing a given set of types could include different neighbouring genomes.

Of 75,774 skani comparisons between the 173-genome hoffmannii lineage and the 438-genome oharae/steigerwaltii/xiangfangensis set, 74,445 met ANI ≥95% and AF ≥80% for both inputs. All nine cross-set type ANIm comparisons gave 95.919–96.020% identity with minimum AF ≥84.781%. The three hoffmannii-associated types shared 99.09–99.12% skani ANI. Figure S2 shows the population comparisons, and Figure S3 shows the within- and between-subspecies ANI ranges for the 437 genomes with assigned oharae, steigerwaltii or xiangfangensis subspecies; the species-only peripheral genome is excluded from Figure S3.

##### *E. cloacae* and *E. dissolvens*

The type-strain comparison is shown in Figure S4.

The comparison contained 205 cross-group pairs, including 164 study–study pairs. All had ANIm 95.287–95.592%, and 200 met bilateral AF ≥80%. Together with the shared grouping of the two lineages in all three phylogenies, these measurements support their treatment as one species with two subspecies. Five comparisons involving CZ862 failed reciprocal AF despite retention of its taxonomic assignment. Reciprocal chromosome ANIb between the types was 95.17022/95.16711%.

##### *E. asburiae* and *E. dykesii*

The type-strain comparison is shown in Figure S5.

All 69 genomes intermediate between the type-associated groups were closer to *E. asburiae* by small ANI and tree-path margins, whereas quartet topology grouped them with *E. dykesii*. These measures describe different aspects of relatedness. We treat the two named groups as one broad *E. asburiae* species; the two outside genomes remain species-unassigned with distinct AF qualifications. A corrected composition parser did not alter the underlying tree result.

##### *E. bugandensis* and *E. nematophilus*

The type-strain comparison is shown in Figure S9.

We retained E. bugandensis and E. nematophilus as separate species because genomic similarity and phylogenetic analyses consistently distinguished two groups. Expanded sampling included 26 bugandensis-associated and six nematophilus-associated genomes. Within-group skani ANI was 97.69–100% and 98.11–99.83%, respectively, compared with 94.69–95.73% between groups. Both groups remained intact across ten local trees on the same 38-genome panel.

This separation persisted despite type-strain ANIm of 95.581% and 138/156 between-group comparisons meeting the ANI ≥95% and bilateral alignment-coverage ≥80% criteria, showing that these thresholds alone did not distinguish the species. Boundary genomes ECC-219 and strain 93 remained unassigned.

Recombination masking removed approximately 55% of the strict-core alignment, but Gubbins reached its iteration limit [47]; subsequent topology tests were therefore considered exploratory. No additional dDDH analysis was performed. The retained species boundaries reflect the combined evidence from the sampled genomes rather than a universal ANI or phylogenetic-distance threshold.

##### *E. chengduensis* and *E. rongchengensis*

The type-strain comparison is shown in Figure S8.

The five-genome comparison included three chengduensis-associated and two rongchengensis-associated genomes, one of which was an additional draft type. Type ANIm was 95.733%, with minimum AF 78.88%, and all six cross-group skani comparisons failed the bilateral AF screen. Although the rongchengensis type has 89 contigs, coverage failures also occurred between complete chromosome inputs, so fragmentation alone does not explain the result. Table 1 of He et al. [33] reports 95.58% ANI and 64.4% dDDH between strain 170250 (E. rongchengensis) and E. chengduensis. We propose subspecies status based on the combined genomic evidence, while acknowledging that the alignment-coverage failures and published dDDH value support retaining separate species.

##### *E. roggenkampii* and *E. quasiroggenkampii*

The type and population comparisons support the proposed quasiroggenkampii subspecies within *E. roggenkampii* (main Table 2; Figure S7). This treatment retains the published dDDH value of 64.8%, which supported separate species in the earlier analysis. The adjacent pseudoroggenkampii/vonholyi comparison below concerns a different boundary and is not resolved by this subspecies proposal.

##### *E. mori* / *E. quasimori* and *E. sichuanensis* / *E. chinensis*

The type ANI and tree relationships support treating *E. quasimori* as a later heterotypic synonym of *E. mori* (main Table 2; Figure S6). *E. sichuanensis* and *E. chinensis* remain separate: their type ANIm was 92.484–92.485%, despite the close relationship recovered by the distance trees. Supplement S04 examines the disagreement with the marker-tree placement and the unresolved culture identity of the alternative *E. sichuanensis* assembly.

##### Other boundaries and unnamed lineages

The *E. pseudoroggenkampii* and historical vonholyi-associated cores contained five and seven genomes, respectively; ECC-249 remained unassigned. CZ-1 was retained within the proposed, non-validly published *E. adelaidei* concept with a boundary qualification. Two ANIm comparisons involving BS0013 addressed a representative gap but did not characterise the entire nine-member GTDB cluster. The BS0013 assembly has 45 contigs and reported depth 2.2×, without a CheckM estimate. The soli-related isolateB comparison included two diagnostic additions, one a 9× metagenome-assembled genome, and one pair failed AF at 79.20%. These observations support lineage comparisons but do not establish a new formal species. CP102 was assigned to *E. soli* at species level only, with its divergent position retained.

RHB15-C17 was retained as *Lelliottia nimipressuralis* context. The two genomes labelled cloacae_P in GTDB and ECC-175 remained unnamed. RHBSTW-00994 was treated as an *Enterobacter* lineage, with uncertainty about genus placement in the broader trees.

#### S07. Genomovar definitions and multilocus sequence typing

The genomovars were defined using 172 study genomes and one additional type-strain genome, giving 173 genomes in total. All 14,878 pairs had AF ≥80% in both directions. We applied complete-linkage clustering to 100 minus ANI using SciPy 1.18.1 and deterministic accession ordering, requiring every pair within a group to reach 99.5% ANI. The threshold was informed by the reported 99.2–99.8% within-species ANI gap [31]; the sampled Enterobacter groups were also checked directly for threshold sensitivity. Fastbaps populations and multilocus sequence types were assessed separately [48, 49].

Sequence types were assigned with mlst 2.32.2 using the seven-locus PubMLST Enterobacter scheme (ecloacae: dnaA, fusA, gyrB, leuS, pyrG, rplB and rpoB). The frozen resource contained 4,751 alleles and 3,291 profiles, restricted to records submitted before 2025; this does not reconstruct an exact 2024 database snapshot. Alleles were identified by BLAST and STs assigned only to exact seven-allele profiles; inexact profiles remained unassigned. STs were supplementary descriptors and did not determine taxonomic assignments [49].

Complete-linkage clustering at inclusive ANI 99.5% partitioned the 173 genomes of the proposed *E. xiangfangensis* subsp. *hoffmannii* into 25 genomovars. Ten groups had multiple sampled members and fifteen were singletons; the largest contained 92 genomes, comprising ninety ST78, one ST2480 and one without an assigned ST. Every within-group pair met the threshold. Eleven pairs from different groups also met or exceeded it, showing that the groups are not separated by a universal pairwise gap (Tables S10 and S11).

Average linkage reproduced the partition. Single linkage produced 23 groups and included twenty-one within-group pairs below the threshold. Changing the threshold to 99.495 or 99.505% did not alter the complete-linkage partition. These sensitivities describe this sampled collection and do not validate a population boundary for future genomes. The seven fastbaps populations are a separate partition and were not treated as equivalent to the genomovars.

Figure S10 shows all saved pairwise ANI values ordered by the unchanged genomovar memberships, together with each group size. Complete linkage requires all within-group pairs to meet the cutoff; it does not require all between-group pairs to fall below it. The eleven cross-group pairs at or above the cutoff are marked in the heatmap and listed in Table S11. The partition is therefore an explicit grouping convention rather than a set of perfectly separated pairwise ANI bands.

#### S08. Reference construction and classifier evaluation

##### Reference selection and study queries

The starting set for classifier development comprised 1,070 study genomes and sixteen additional types. To limit closely related records crossing the development and internal-reserve partitions, genomes were linked using shared BioSamples, adjudicated assembly identity, identical whole-file hashes and chromosome skani similarity ≥99.9% ANI with both AF values ≥90%. Shared BioProjects also linked records for partition allocation. Type-containing blocks and representation of eligible species/subspecies strata were retained in development. Other blocks were allocated using a deterministic SHA256 rule with seed 20261004 and nominal reserve probability 20%. This produced 1,034 development genomes and 52 internal-reserve genomes. All had contributed to the broader taxonomic study, so the reserve was not an untouched external test.

The 140 primary references retained all 33 historical type-strain accessions and sampled up to eight genomes per reviewed species/subspecies stratum. Where no type was present, the genome with the smallest total chromosome-ANI distance to the other genomes in its development stratum (the medoid) was selected first. Subsequent selections maximised the distance to the closest already selected reference, with deterministic quality tie-breaking. Supplemental genomes required recorded NCBI completeness ≥98% and contamination ≤2%; three types falling below the supplemental completeness requirement were retained explicitly. The panel covered 31 sampling strata, 24 species concepts and ten subspecies concepts (Table S12). Whole assemblies were used for this classifier reference component. A second component retained all 173 chromosome sequence sets that define the genomovars, because matching a genomovar requires comparison with every member, not only a representative.

The primary panel contained 124 study genomes and all sixteen additional types. Removing these references from the 1,070 study genomes left 946 study queries: 922 with named-species assignments and 24 unnamed or contextual genomes. Their classification measured reproduction of the study labels. Of the 922 named genomes, 836 had individually reviewed taxonomic treatments and 86 retained compatible source names. There were 107 near-clones of references; selecting one genome per clone block left 410 named genomes. This reduces repeated sampling of related sequences but does not make the labels independent of the study.

##### Assignment rules in Pterobacter

Species assignment requires skani ANI ≥96% and AF ≥80% for both genomes, met by the same reference in the nearest species, together with a measured comparison to an alternative species. Subspecies assignment additionally requires ANI ≥98%, the same coverage criteria and a margin of at least one percentage point over competing subgroups within the species. Ambiguous matches and identified reference or sequence-scope problems leave the affected rank unassigned. These conservative identification rules differ from the 95% ANI screen used to investigate taxonomic relationships.

The Rust implementation adds alignment-based comparison when a query’s maximum returned skani ANI is ≥94.5% but <96%. Assignment through this route requires a unique nearest named species, a reference meeting both skani coverage criteria, reciprocal ANIm strictly >96% and all four directional ANIm coverage measurements ≥80%. The same reference must meet the identity and coverage requirements. MUMmer4 4.0.1 was used for these reciprocal alignments [50]. Before assigning a species, the tool checks the active reference panel for competing matches; missing evidence, an incompatible match or a disallowed disagreement between ANI methods prevents assignment. ANIm cannot promote a subspecies or genomovar assignment. The following sections define the evaluation, and Supplement S10 records the evaluated build and numerical procedure.

Genomovar identification requires an eligible assignment to *E. xiangfangensis* subsp. *hoffmannii*. The query must then meet ANI ≥99.50% and bilateral AF ≥80% against every member of one reference group. Each alternative group must be excluded by a measured comparison below the threshold. Matches to more than one group remain unresolved; missing comparisons cannot exclude a rival. Matches to a group represented by a single genome carry a warning about limited reference coverage.

##### Additional public genomes and review of disagreements

The eligible public metadata pool contained 6,424 assemblies after exclusions for known earlier development, reference and diagnostic links. Records were grouped conservatively using accession lineages, explicit GenBank/RefSeq pairings, BioSamples, BioProjects and exact normalised strain labels. Such links control known dependence without proving isolate identity. A metadata-selected set of 96 assemblies was used for method confirmation, and 1,471 additional members of their linked groups were held out, leaving 4,857 assemblies for evaluation. These were non-complete assemblies: 3,551 were deposited at contig level, 1,293 at scaffold level and thirteen at chromosome level. Thirty-six of these had subsequently been exposed during development and were kept separate from the 4,821-assembly source-name comparison. Figure 1 shows this selection. The final comparison set contained 788 linked components.

Public names were mapped to the study’s taxonomic concepts before comparison. Of the 4,821 assemblies, 4,387 had a species expectation that could be scored; 434 had descriptive outcomes only. Using one genome per linked component, 663 of 687 scored components agreed with the mapped species name, eleven differed and thirteen were unassigned. These comparisons measure agreement with deposited names, not independent accuracy.After classification, 95 distinct species/subspecies disagreements were found in the source-name comparison. After excluding four genomes already reviewed, a fixed hash-based stratified draw selected twenty of the remaining 91 for review using ANI and marker and distance trees with the coded query genomes added. These twenty genomes were selected for review because Pterobacter disagreed with their deposited names. Examining a broader set of genomes in the phylogenetic trees changed the confidence ratings for two assignments, but did not change any taxonomic labels. All twenty final assignments were rated as high confidence. All twenty reviews agreed with the classifier’s species assignment, and all nineteen with an explicit subspecies assignment agreed at that rank. Because these cases were selected following disagreement with deposited names, they were not an independent accuracy sample and do not establish outcomes for the other 71 eligible disagreements.

##### Reference-removal settings and alignment-assisted classification

We tested how reference availability affected classification by evaluating 1,034 development genomes under nine settings. Three levels of removal excluded the query and its aliases, its near-identical genomes, or its broader group linked by sample or project records. Each was combined with retaining the expected taxon, removing its species, or removing its subspecies where applicable. This produced 9,306 classification outcomes from repeated tests of the same genomes. Overall, when references from the expected subspecies were missing, Pterobacter could still assign some queries to the correct species using ANIm, whereas removing all references from the expected species yielded no additional ANIm-assisted assignments.

The ANIm-assisted classifier was also evaluated on the original study and public datasets and eleven additional genomes near a species boundary. Six of these eleven had labels previously informed by ANIm and were considered separately from the other five.

ANIm was considered when the highest skani ANI was ≥94.5% but <96%. A qualifying reference had to meet bilateral skani coverage ≥80%, ANIm >96% in both alignment directions, and coverage ≥80% for both genomes in each direction. Final checks withheld assignment if there was a competing species match, disallowed disagreement between ANI methods, or insufficient evidence.

Across all evaluation datasets and settings, 2,740 classification attempts triggered ANIm evaluation, of which 2,562 remained unassigned because no reference satisfied all the required identity and coverage criteria. 178 had at least one qualifying reference: 123 passed the final checks and gained a species assignment, while 55 remained unassigned because of ANI-method consistency checks. These counts describe repeated classification attempts, not distinct genomes or assignments lost after reference removal.

All 123 additional assignments occurred when the expected subspecies was removed: 50 with query/alias removal, 50 with near-clone removal and 23 with broader linked-group removal. ANIm added no assignments in the other settings or datasets, including the original study and public-genome runs. None of the eleven additional boundary genomes had a qualifying reciprocal ANIm match.

No competing-species match was found in the 178 complete-panel checks. However, these checks did not test cases where skani failed to return a relevant reference. Ten incomplete outcomes involved repeated tests of one query lacking a comparison with an alternative species, rather than alignment failures. The experiments therefore describe behaviour under the tested reference conditions, rather than an independent improvement in classification accuracy. Figure S11 summarises the reference-removal outcomes, and Tables S13 and S14 provide detailed classification and reference-comparison results; Supplement S10 records the implementation.

##### Genomovar evaluation

We evaluated genomovar assignment by removing individual queries, related genomes or entire genomovars from the reference set, and by testing additional public assemblies and artificially fragmented genome inputs. The checks assessed whether the software followed its assignment rules, rather than independently establishing biological accuracy. Overall, genomovar assignment depended on reference coverage: removing individual queries usually preserved their original group assignments, whereas removing related genomes often prevented assignment and removing an entire genomovar could produce misleading matches to neighbouring groups.

Removing each query from its definition group gave 148 source-group matches, ten ambiguous matches and fifteen unassessable singletons among 173 genomes. Removing related genomes more broadly gave 26 source-group matches, 123 incomplete results, nine ambiguous matches and fifteen unassessable singletons. Among 749 public assemblies assigned to the parent lineage, 539 uniquely matched a multi-member group, 26 matched a singleton, 71 matched overlapping groups, two failed member coverage and 111 matched no group. These datasets are identified in the source tables as GV1b, GV2 and GV3-public, respectively.

To test the effect of an absent group, all references from a query’s own genomovar were removed. Ten of 173 definition genomes then matched a neighbouring group: nine GV05 genomes matched GV04 and one GV01 genome matched GV24. Two of 322 fragmented derivatives, each 2 kb long, showed matches in the reverse direction. All twelve had been ambiguous with the complete reference panel. These matches were recorded as diagnostics and were not emitted as deployment assignments. Figure S12 illustrates this effect, and Table S15 lists every case. The record-level exact intervals are 2.81–10.37% for ten of 173 and 0.08–2.23% for two of 322; correlated derivatives and shared definitions prevent interpreting them as population accuracy estimates.

Of the 322 fragmented genome derivatives, 151 received the prerequisite species and subspecies assignments and proceeded to genomovar matching. The remaining 171 stopped before this step because their species or subspecies assignment was unresolved or ineligible. Across the wider implementation checks, 670 complete genomovar-rule replays and these 171 parent stops gave 841 cases. The records also account for 914 native genomovar contexts and thirty saved controls, comprising twenty-five classifications and five expected refusals for queries outside the applicable lineage.

#### S09. Kraken classification with a general-purpose database

Kraken2 2.1.6 with Standard 20260226 was run at confidence 0 and 0.1 on both whole-assembly contigs and deterministic non-overlapping fragments of at most 150 bp, including terminal fragments. Both representations contained the same 5,382,995,503 bases from 1,070 assemblies. Each sequence’s reported taxon was weighted by its entire length; unclassified sequences remained in the denominator. This is a sequence-call-length summary, not a measurement of the taxonomic support at each base.

Increasing Kraken’s confidence threshold from 0 to 0.1 reduced the proportion of assembled sequence assigned to species level or below from 99.961% to 16.502%; the remaining sequence received broader taxonomic assignments or remained unclassified. The corresponding fragment values were 25.722 and 19.021%. Table S16 supplies integer sequence and base totals and the denominator for each percentage. The artificial fragments describe sequence-length sensitivity and do not estimate performance on empirical reads.

Figure 3 compares Kraken’s names with the revised assignments for the hormaechei-related lineages; Figure S13 extends the comparison to all 32 assignment categories. Strain labels were collapsed only to their recorded species or subspecies ancestor in the saved Kraken taxonomy. Each row is normalised by its own represented input length. The full cross-tabulation retains every call; infrequent columns are combined for display only. These figures compare names without imposing a new species concept on names whose original circumscription is uncertain. Figure S14 retains the four-setting resolution comparison.

Using the original assembly contigs at Kraken’s default confidence setting, 37.073% of the total sequence length received subspecies assignments incompatible with our revised classification, and a further 0.387% received conflicting species assignments. Some results could not be compared directly because the reference database and our study used different species boundaries. These included classifications involving E. chengduensis and E. rongchengensis, which were excluded from the conflict totals. The percentages therefore describe directly comparable naming disagreements, rather than all disagreements or the proportion of incorrectly identified isolates.

Accession-based checks detected representation of 912 of the 1,070 study assemblies in the Kraken2 reference database. No match was detected for the remaining 158, but their absence from the database could not be established.

One sequence-level example illustrates the confidence effect: AP022126.1 from the *E. kobei* assembly GCA_014169295.1 was assigned to an *E. cloacae* strain at confidence 0 and to the genus Enterobacter at confidence 0.1. Different sequences from an assembly can receive different labels; neither this example nor the length-weighted summaries should be interpreted as a count of misidentified genomes.

#### S10. Evaluated software and computational measurements

The evaluated Rust source was commit 834f4c288e9c163f316f3668f0219004d0adc6da, before adoption of the Pterobacter name; its executable reported terobacter 0.0.0. The embedded skani profile was skani-0.3.2-avx2-compatible-dna-v1. The initial Python classifier assessment used version 0.2.0. These evaluated versions are distinguished from subsequent release candidates. Primary references comprised 140 whole assemblies, while the genomovar component retained 173 chromosome definition sets. Whole query assemblies were not trimmed to improve AF. Reciprocal nucleotide alignment requires sequence data in addition to sketches.

The alignment-assisted evaluation used MUMmer4 4.0.1, reciprocal orientation, N-preserving input and the retained MUM/one-to-one span procedure, identified as reciprocal-mummer4-4.0.1-preserve-n-mum-1-span-v1. Classification requested 11,790 distinct query–reference pair keys; the 11,791 retained pairs included one additional parser-check pair. Thirty-three pairs for the eleven additional boundary queries were computed after those labels were fixed, although six labels had already been informed by earlier ANIm evidence. Fifteen integrated runs used default parameters and three used explicitly supplied default-valued custom parameters. Timings for individual alignment workers were kept separate from end-user execution time.

Tables S17–S19 report computational measurements for 5,803 completed native queries, 11,791 reciprocal pairs and eighteen integrated runs, respectively. For the initial classification rules, median elapsed and CPU time were 1.72 and 2.25 s, with median peak resident memory 226,492,416 bytes. Pairwise alignment had median summed native-worker CPU time 29.34 s and median per-pair maximum native-worker resident memory 129,679,360 bytes; these measures exclude controller, archive and scheduler overhead. The fifteen default integrated runs had median elapsed time 38.14 s and median peak resident memory 191,614,976 bytes. These measurements describe the evaluated Linux configuration. The complete Verticall phylogenetic calculation used an estimated 2,784.8 allocated core-hours, which is a resource-allocation measure rather than process CPU time.

Software fixtures tested classification rules, numerical input handling and simulated failures separately from the biological datasets. They are not additional biological samples. A subsequent platform comparison used sixty public assemblies, 120 skani component comparisons, forty-four reciprocal alignment pairs and classification under two policies. Following correction of an alignment-filter tie-ordering difference, the results matched exactly between Linux and an Apple M4 Pro running macOS 15.7.7. That comparison used source commit c29178f4f2a8383d39917eb9582aac3627642828, separate from the biological evaluation above. It establishes reproducibility on the fixed comparison panel, not additional biological accuracy. Pterobacter 0.1.0 was subsequently released at https://github.com/mbhall88/pterobacter/releases/tag/0.1.0. Source versions, reference hashes, retained commands and computational records are provided through the analysis repository and archive identified in the main Data and code availability statement; the repository documents missing historical commands and limits of end-to-end reconstruction.

#### S11. Nomenclatural evidence and limitations

Tables S20, S21, S22 and S23 list the proposed name changes and their nomenclatural status, identify the type-strain culture deposits and assemblies, show which previously published names had type genomes included in the priority check, and give genome-size and G+C summaries, respectively. Whole deposited assemblies supplied genome sizes and taxon ranges. G+C contents were calculated from integer A, C, G and T counts. These descriptions use the original published phenotypic properties; no new phenotyping or dDDH measurement was performed.

LPSN records were retrieved on 8 October 2026. To check whether an earlier published name could take priority over a proposed name, the screen considered 53 validly published species names and selected type assemblies for 49. *E. nimipressuralis* had NCBI type-material candidates but could not be resolved by the applied alias matching; it was not assumed to lack a genome. *E. pulveris* and *E. pyrinus* are historical combinations now placed outside *Enterobacter*. *E. siamensis* remained unresolved. Priority based on page order addresses the xiangfangensis name but does not exclude every possible competitor among merged taxa or later subspecies epithets. Forty-seven GTDB R232 matches used the selected assembly version, whereas *E. massiliensis* and *E. sacchari* matched a different version. These are metadata and coverage limitations, not new phylogenetic results.

Five transferred combinations and three corresponding type-subspecies combinations are proposed in the main text (summarised in Table 4), together with emendations of *E. xiangfangensis* and *E. hormaechei*. Nomenclatural dates distinguish effective publication from validation: the oharae and steigerwaltii names were validated in 2016, hoffmannii in its earlier combination in 2018, rongchengensis and intestinihominis in 2025, and the hoffmannii/nototheniae names in 2026. The Sedláček et al. description belongs to Czech Polar Reports 15(2), 2025, despite its 2026 online date; the date of nomenclatural standing follows Validation List 230.

The proposed quasiroggenkampii and rongchengensis subspecies treatments retain the published dDDH values of 64.8 and 64.4%, respectively, and the associated coverage limitations (main Table 2). Their interpretation differs from retaining separate bugandensis/nematophilus cores despite ANI overlap. These are evidence-based taxonomic treatments of the sampled groups, not applications of a newly optimised universal threshold. The proposed *E. adelaidei* name lacks valid publication, and isolateB remains unnamed. The proposed combinations are not yet validly published.

#### S12. Public database and supporting files

The reference database version 2026-10-15 is deposited at https://doi.org/10.6084/m9.figshare.34640883.v1 [42]. Its published archives and content hashes were verified independently of the frozen classification evaluation. This deposit does not change the evaluated software version or any results reported above. The 23 revised supplementary tables accompany this manuscript as individual TSV files and as worksheets in Enterobacter-supplementary-tables.xlsx. The earlier table version is archived at https://doi.org/10.6084/m9.figshare.34742010.v1; its numbering differs from this revision. Analysis scripts and selected source data are available at https://github.com/mbhall88/Enterobacter-analysis (commit 3a2baebaa1c82149451fe6039659bfc83ec1ca5a); the corresponding sequences, alignments, trees and frozen evaluation packages are archived at https://doi.org/10.6084/m9.figshare.34742013.v1.

#### Supplementary figures

Figures S1, S4–S9 show selected type strains and named neighbours pruned from the same three saved 35-tip projections used for main Figure 2. They orient the taxonomic comparisons; they do not replace the population analyses or establish that omitted genomes lack conflicting relationships. These supplementary projections were not re-inferred or rerooted; Figure 2 uses the same source trees with the outgroup display orientation described in S03. Pairwise path lengths among retained tips are preserved. Distance scales are method-specific and should not be compared across panels. Dots retain source SH-like local support (BAC120) or sketch-seed frequency (Mashtree) ≥0.95 for the original branches; support was not recalculated after pruning, and FastME has no branch-support estimates.

The heatmaps use the original skani type–type comparisons of selected chromosome inputs, including all contigs for draft types. Minimum AF is the smaller aligned fraction of the two inputs, not a reciprocal ANIm coverage result. Diagonals are unmeasured self comparisons and are left blank. Colour scales are fixed across all seven figures. Cell labels are rounded to one decimal place; exact values are supplied in the figure source data. Historical type names identify the sequences; current proposed names and full aliases are given in Table S03. Figure S2 adds the population-level ANI evidence for *E. xiangfangensis* and *E. hormaechei*. Standalone PDF, SVG and 300 dpi PNG figures and their exact source tables accompany the supplement.

##### Figure S1. *E. xiangfangensis* and *E. hormaechei*

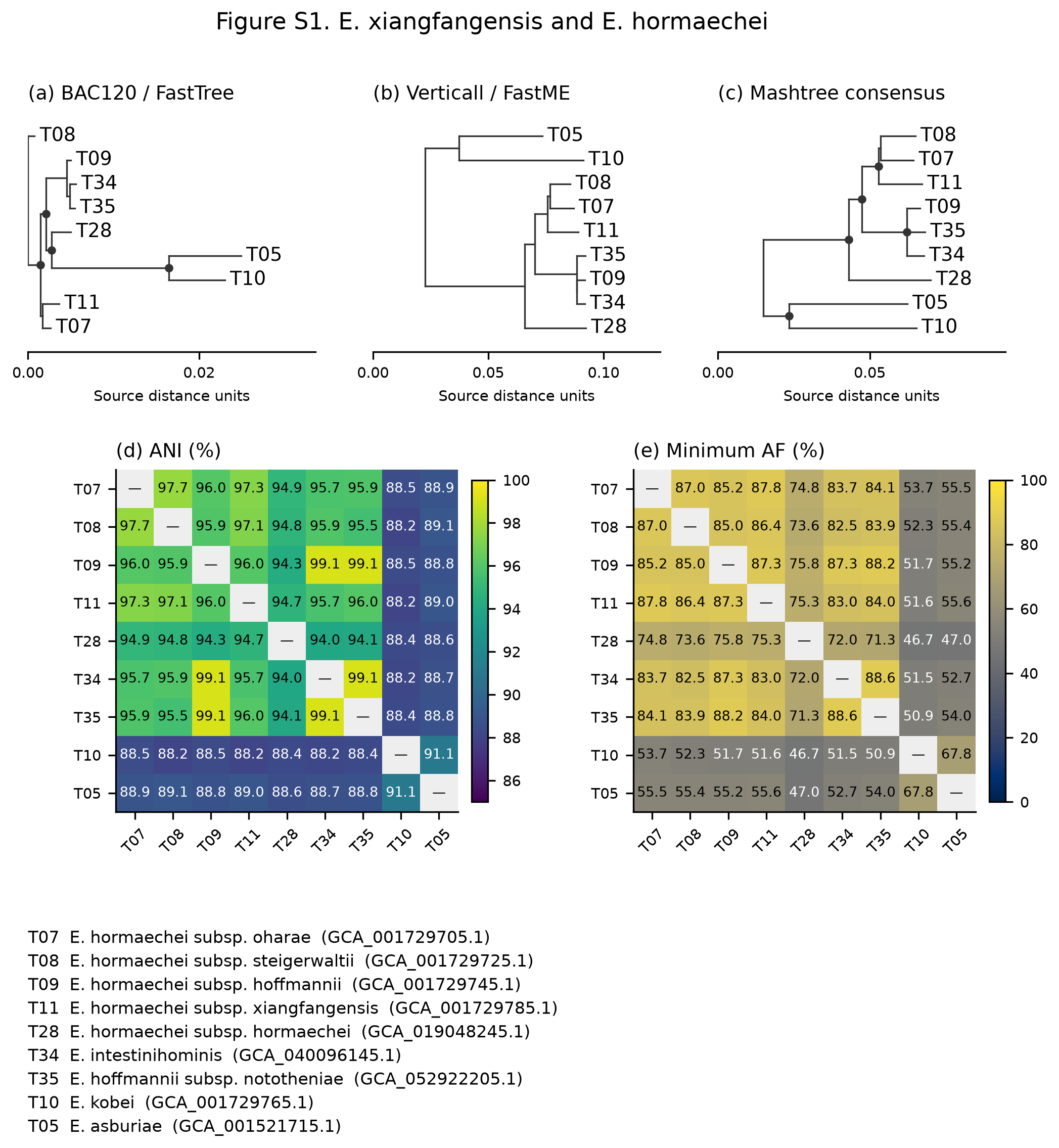

Figure S1

The oharae, steigerwaltii, xiangfangensis, hoffmannii, intestinihominis and nototheniae types are shown with the hormaechei type and kobei/asburiae context. Under the proposed treatment, the first six types belong to *E. xiangfangensis*; the hormaechei type remains in *E. hormaechei*. The three hoffmannii-associated types have an unresolved internal history (S04). Panels (a–c) show the three saved type-tree projections; panels (d–e) show skani ANI and the smaller of the two aligned fractions. Labels identify the historical type names, not automatic classifications of surrounding genomes. The common display rules above apply.

##### Figure S2. Population ANI and alignment coverage

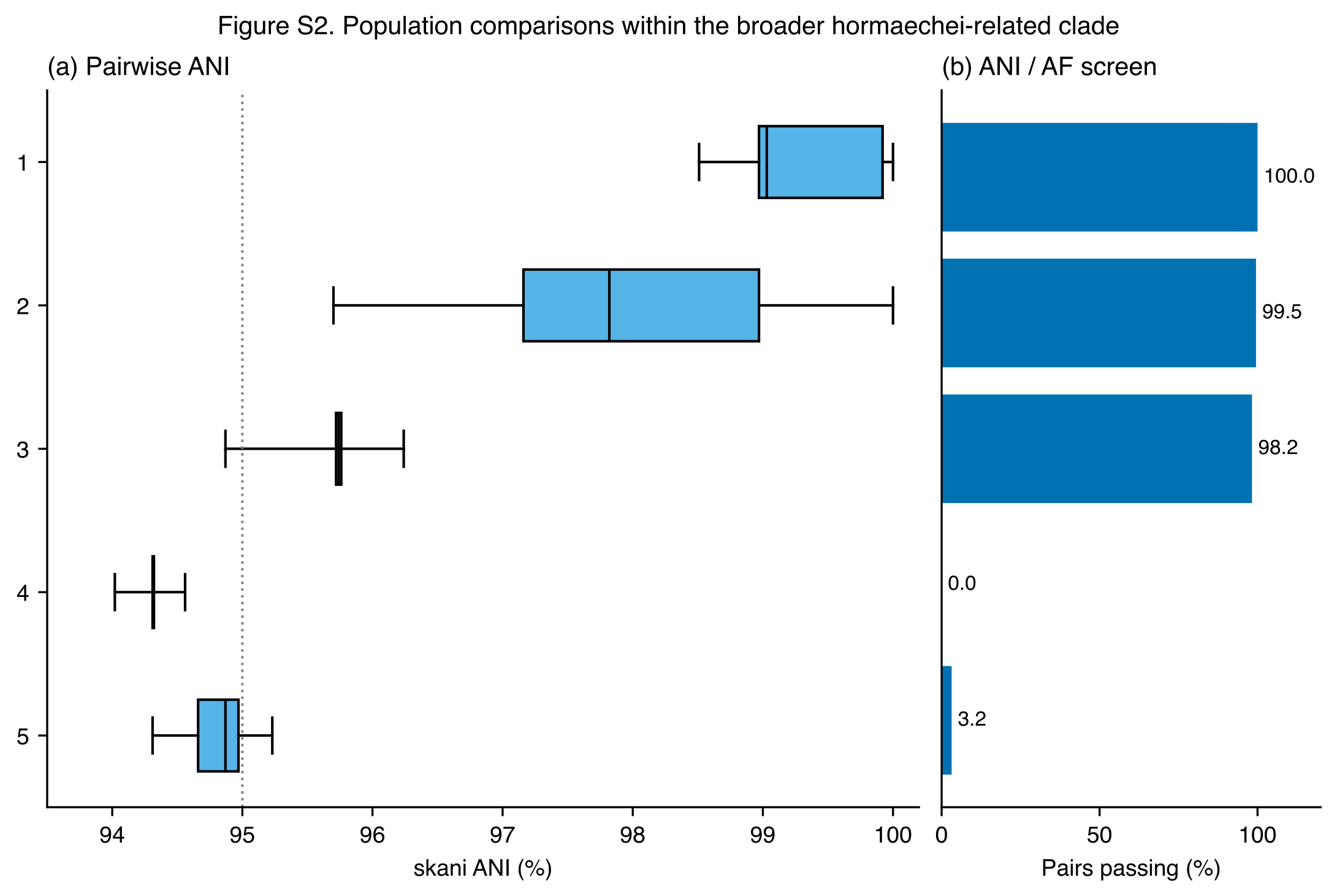

Figure S2

The numbered comparisons are: 1, within hoffmannii lineage (n = 14,878 pairs); 2, within oharae / steigerwaltii / xiangfangensis set (n = 95,703 pairs); 3, between the two *E. xiangfangensis* sets (n = 75,774 pairs); 4, hoffmannii lineage vs *E. hormaechei* (n = 2,249 pairs); 5, other *E. xiangfangensis* set vs *E. hormaechei* (n = 5,694 pairs). The sets contain 173 hoffmannii-lineage genomes, 438 oharae/steigerwaltii/xiangfangensis-associated genomes and thirteen *E. hormaechei* genomes. Boxes show the median and interquartile range; whiskers show the minimum and maximum, not confidence intervals. Panel (b) gives the proportion passing ANI ≥95% and AF ≥80% for both genomes. The 438-genome set includes one peripheral genome assigned only to species. Pairwise observations share genomes and are not independent replicates. These are the saved chromosome-scope skani comparisons, not whole-type or reciprocal ANIm measurements.

##### Figure S3. ANI within and between three *E. xiangfangensis* subspecies

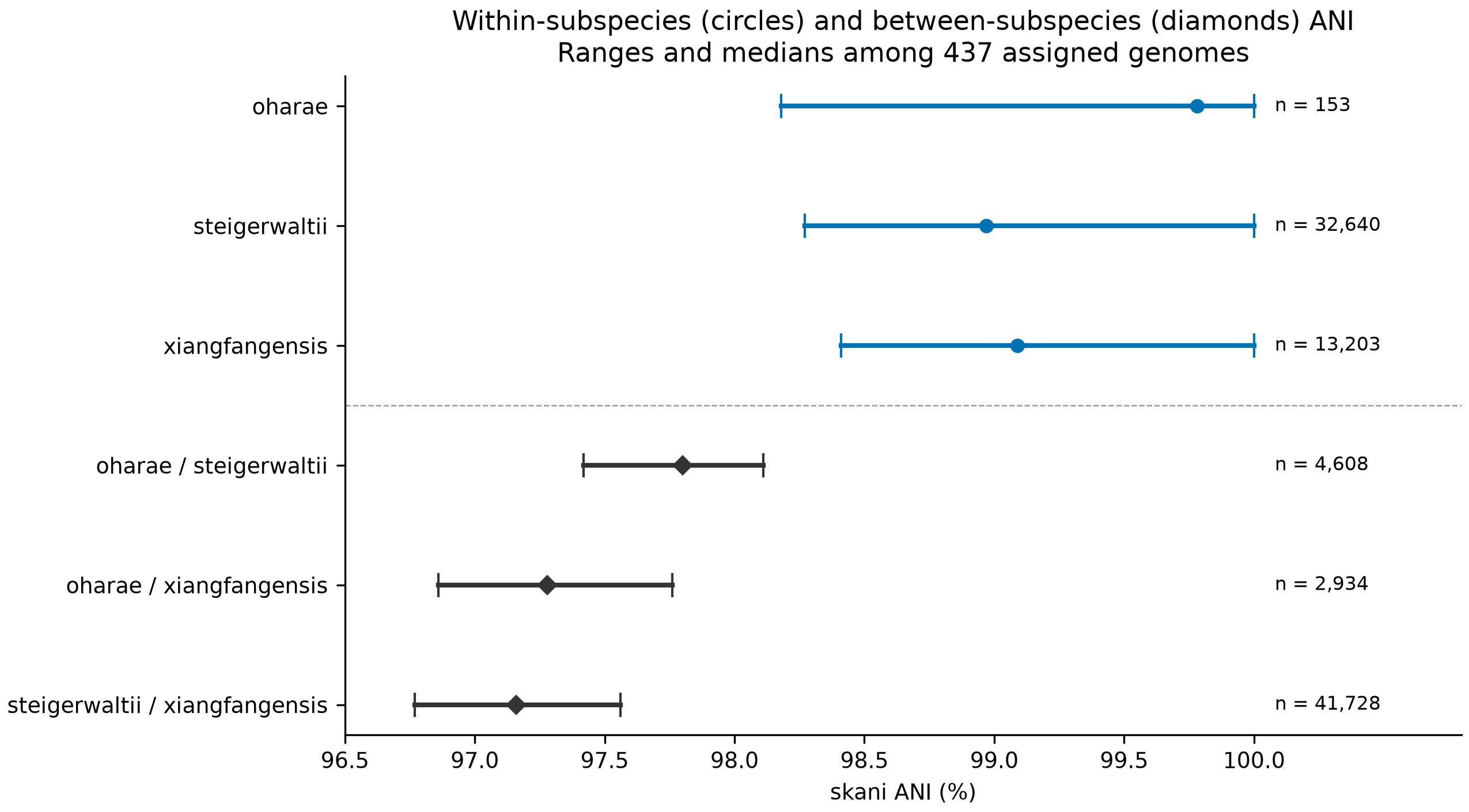

Figure S3. Observed ANI ranges and medians within and between the oharae, steigerwaltii and xiangfangensis subspecies.

The comparisons contain 437 genomes assigned to subsp. oharae (18), subsp. steigerwaltii (256) or subsp. xiangfangensis (163). Horizontal lines show observed minima and maxima; circles and diamonds mark medians for within- and between-subspecies pairs, respectively. Values of n count genome pairs, not independent biological replicates. These are chromosome-scope skani comparisons. Ranges describe sampled variation, not confidence intervals or universal species thresholds.

##### Figure S4. *E. cloacae* and *E. dissolvens*

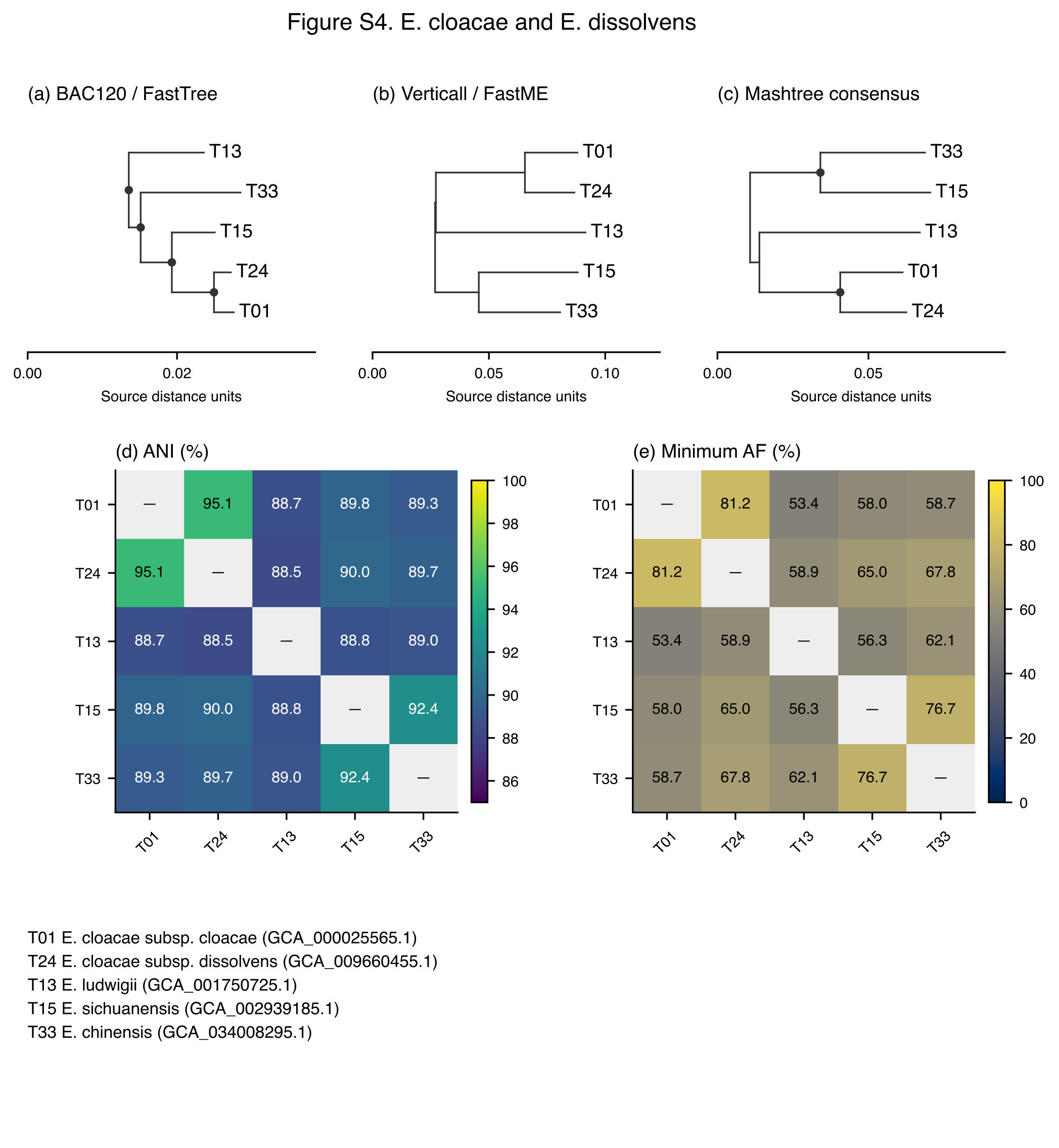

Figure S4

The cloacae and dissolvens types are shown with ludwigii, sichuanensis and chinensis context. The proposed treatment retains two subspecies within *E. cloacae*. The alternative placement of *E. ludwigii* remains unresolved. Panels (a–c) show the three saved type-tree projections; panels (d–e) show skani ANI and the smaller of the two aligned fractions. Labels identify the historical type names, not automatic classifications of surrounding genomes. The common display rules above apply.

##### Figure S5. *E. asburiae* and *E. dykesii*

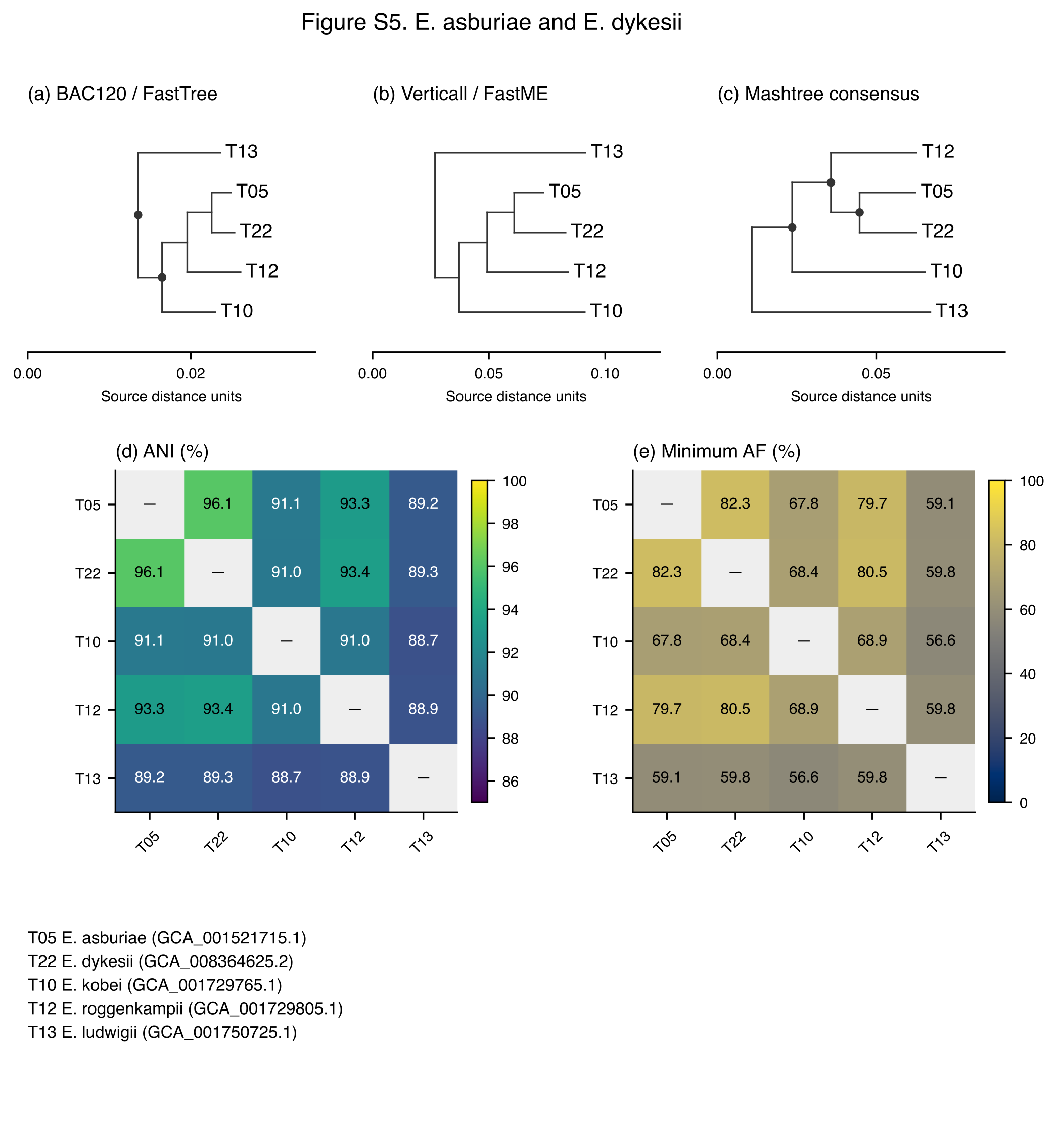

Figure S5

The asburiae and dykesii types are shown with kobei, roggenkampii and ludwigii context. The proposed treatment places *E. dykesii* as a later heterotypic synonym of *E. asburiae*; relationships of the intermediate population are described in S06. Panels (a–c) show the three saved type-tree projections; panels (d–e) show skani ANI and the smaller of the two aligned fractions. Labels identify the historical type names, not automatic classifications of surrounding genomes. The common display rules above apply.

##### Figure S6. *E. mori* / *E. quasimori* and *E. sichuanensis* / *E. chinensis*

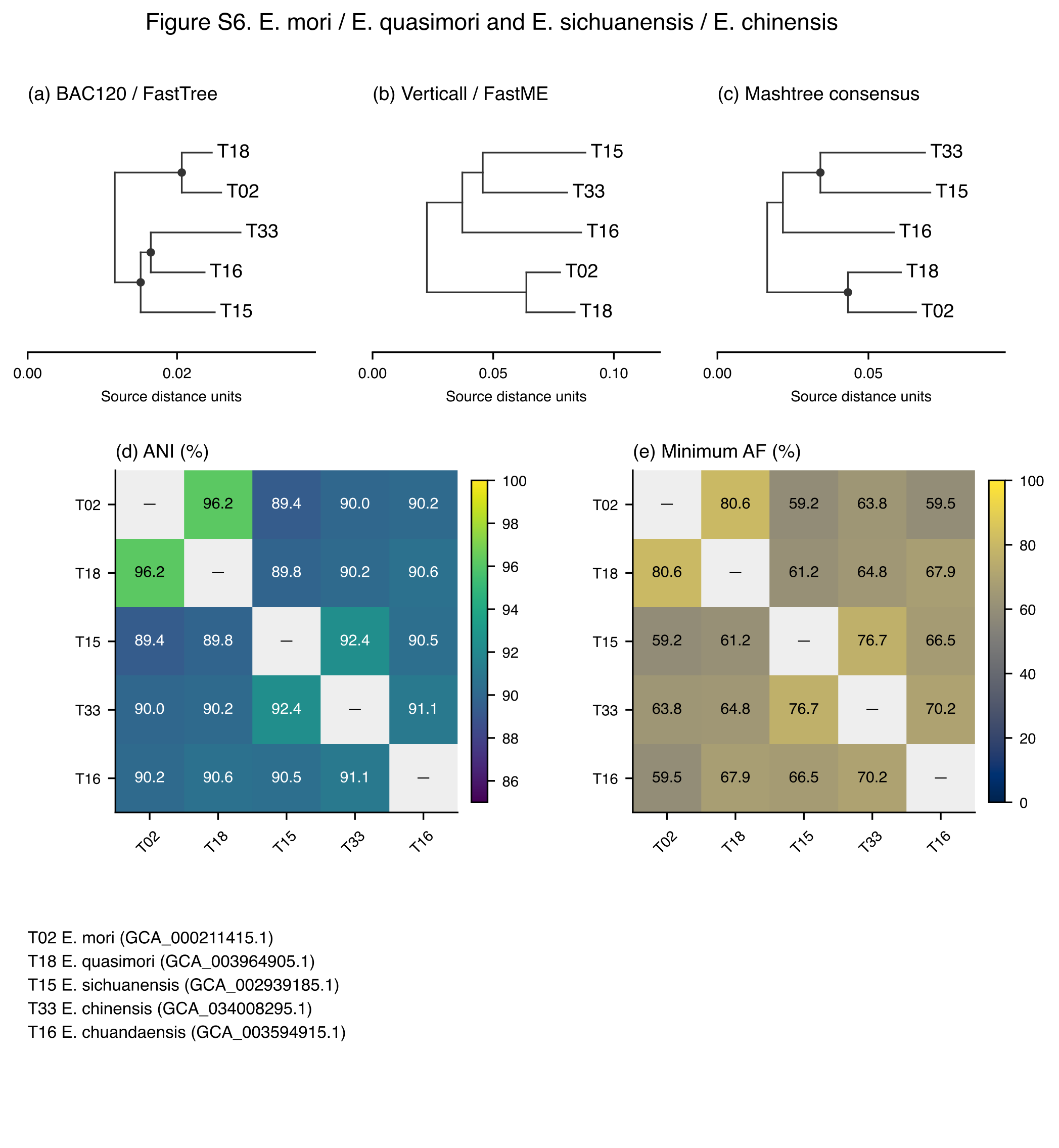

Figure S6

The mori and quasimori types, and the sichuanensis and chinensis types, are shown with chuandaensis context. *E. quasimori* is treated as a later heterotypic synonym of *E. mori*, whereas *E. sichuanensis* and *E. chinensis* remain separate species. Their conflicting tree placements are examined in S04. Panels (a–c) show the three saved type-tree projections; panels (d–e) show skani ANI and the smaller of the two aligned fractions. Labels identify the historical type names, not automatic classifications of surrounding genomes. The common display rules above apply.

##### Figure S7. *E. roggenkampii* and neighbouring types

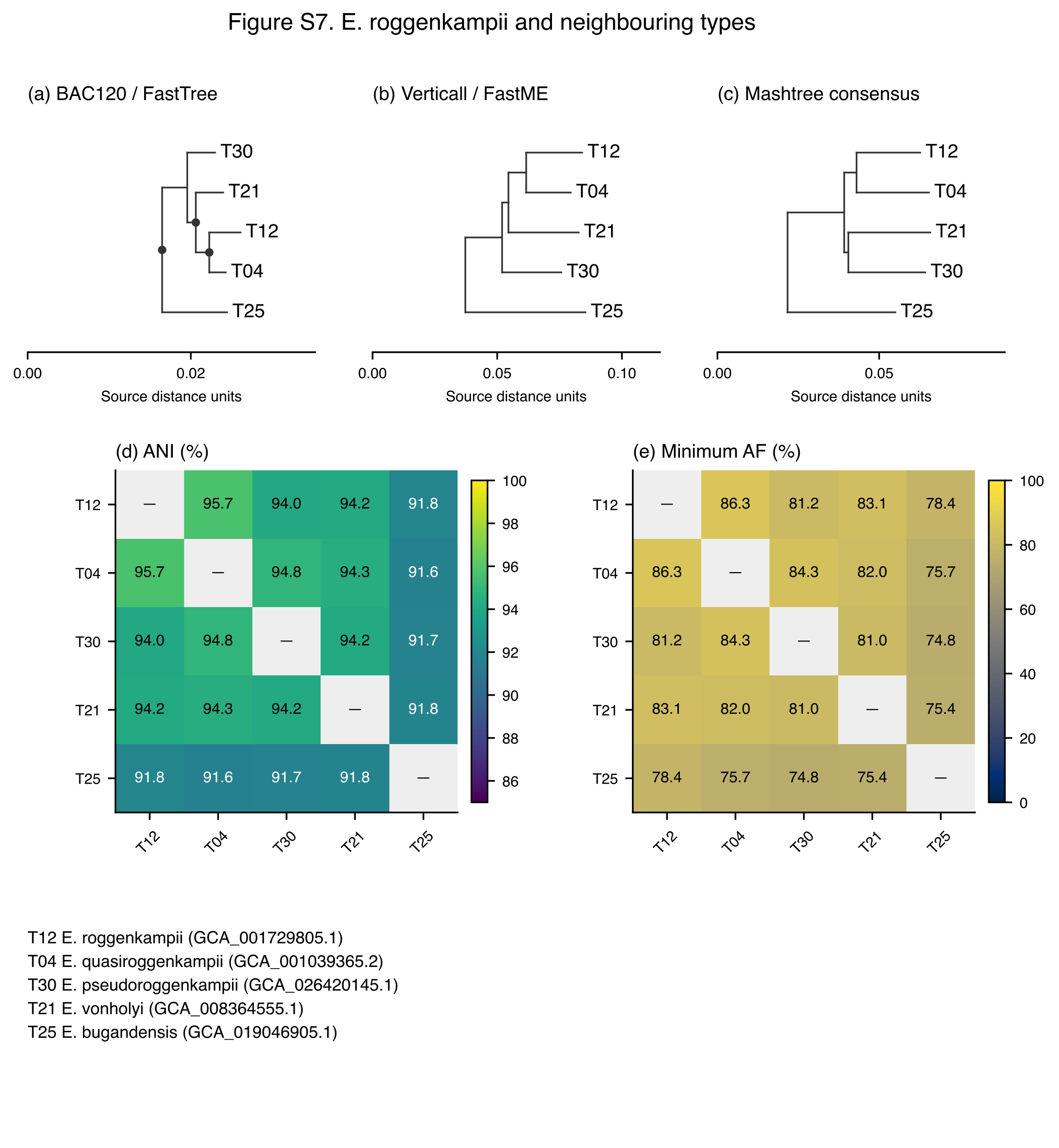

Figure S7

The roggenkampii, quasiroggenkampii, pseudoroggenkampii and vonholyi types are shown with bugandensis context. The proposed quasiroggenkampii subspecies treatment and unresolved pseudoroggenkampii/vonholyi boundary are distinct interpretations; close tree placement alone does not assign the boundary genomes. Panels (a–c) show the three saved type-tree projections; panels (d–e) show skani ANI and the smaller of the two aligned fractions. Labels identify the historical type names, not automatic classifications of surrounding genomes. The common display rules above apply.

##### Figure S8. *E. chengduensis* and *E. rongchengensis*

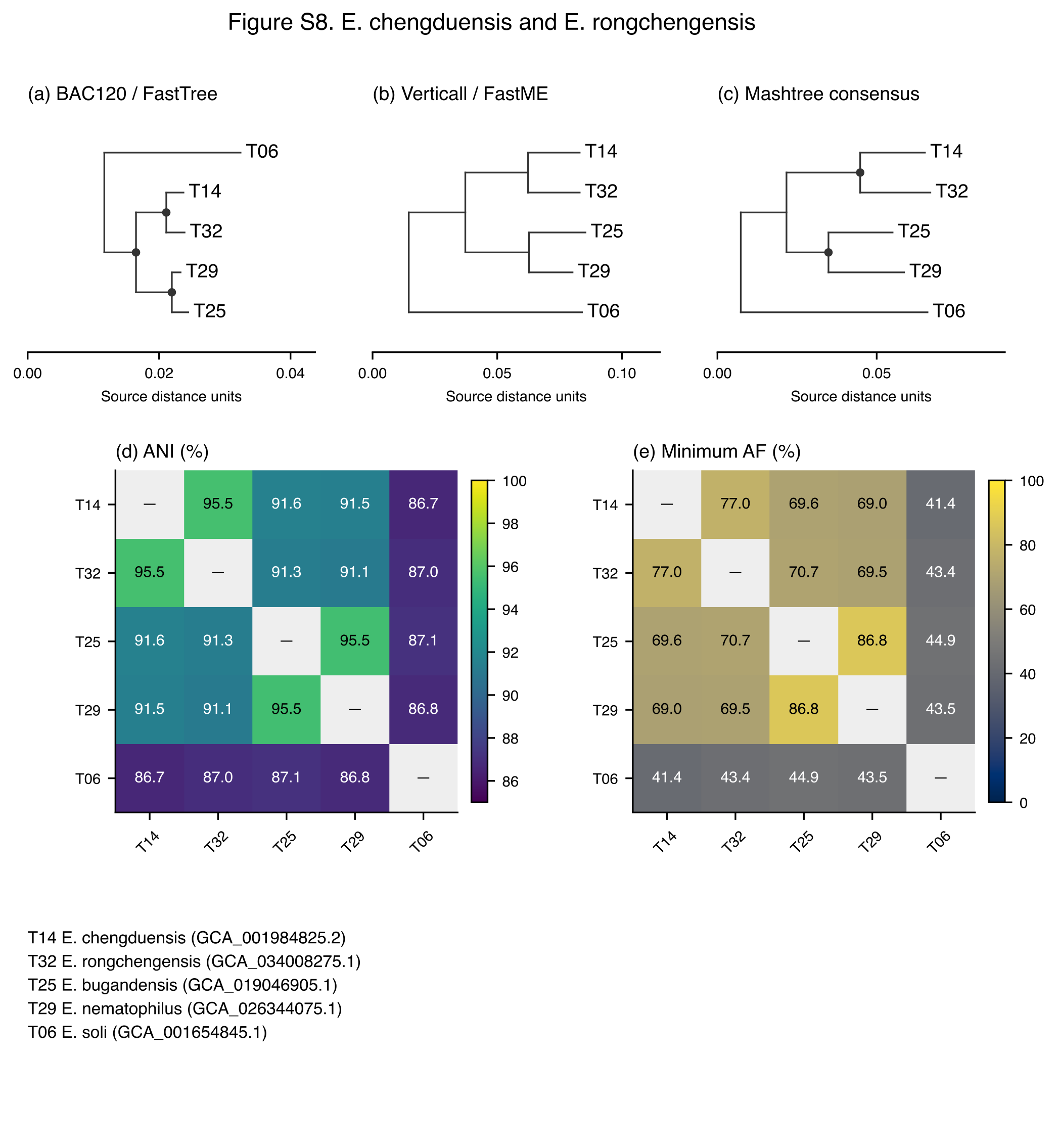

Figure S8

The chengduensis and rongchengensis types are shown with bugandensis, nematophilus and soli context. Their proposed treatment as subspecies retains the conflicting alignment-coverage and published dDDH evidence in S06. Panels (a–c) show the three saved type-tree projections; panels (d–e) show skani ANI and the smaller of the two aligned fractions. Labels identify the historical type names, not automatic classifications of surrounding genomes. The common display rules above apply.

##### Figure S9. *E. bugandensis* and *E. nematophilus*

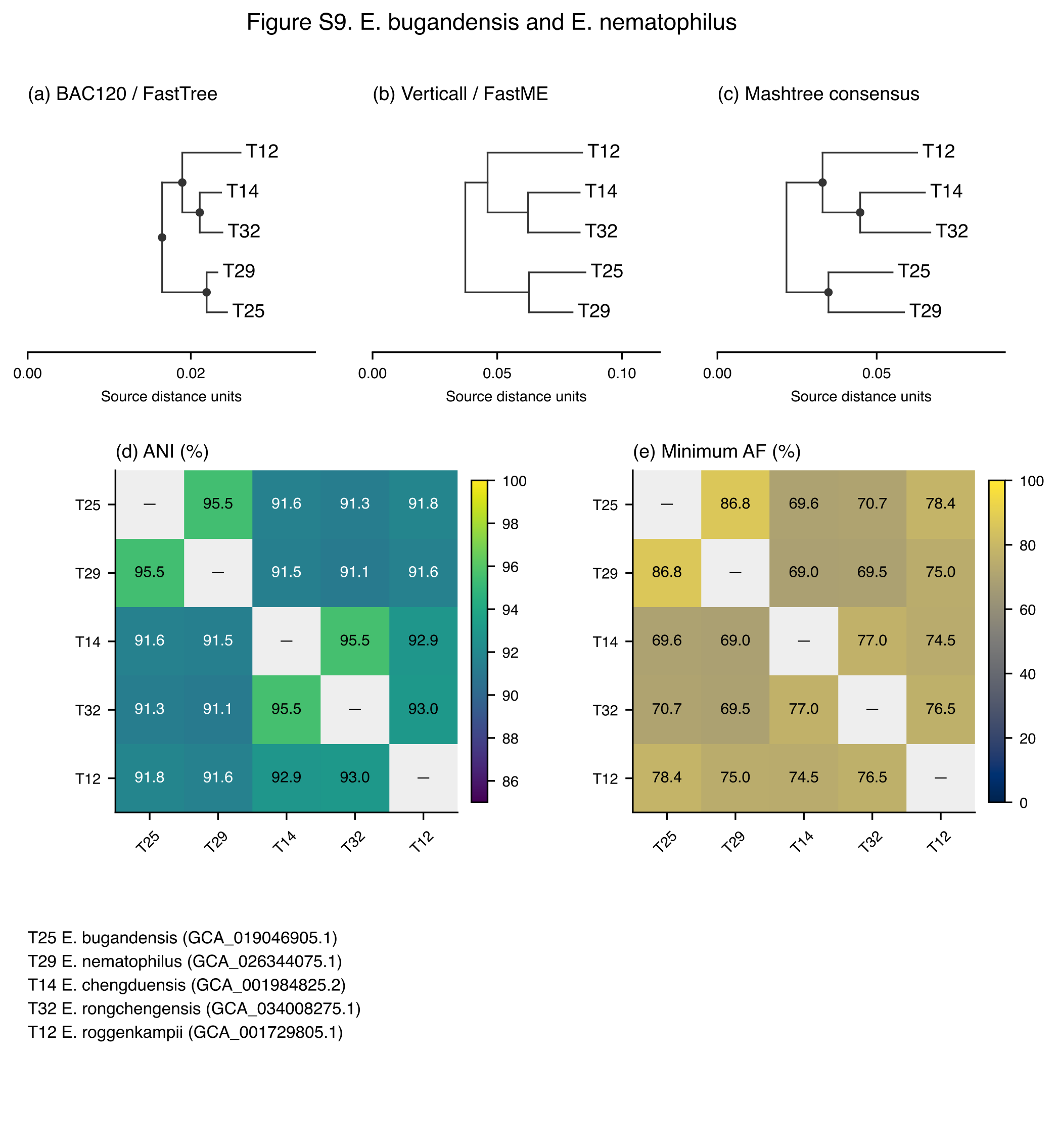

Figure S9

The bugandensis and nematophilus types are shown with chengduensis, rongchengensis and roggenkampii context. The named cores remain separate species despite overlapping ANI evidence. Type-only projections do not show the unassigned boundary genomes. Panels (a–c) show the three saved type-tree projections; panels (d–e) show skani ANI and the smaller of the two aligned fractions. Labels identify the historical type names, not automatic classifications of surrounding genomes. The common display rules above apply.

##### Figure S10. Genomovars within the hoffmannii subspecies

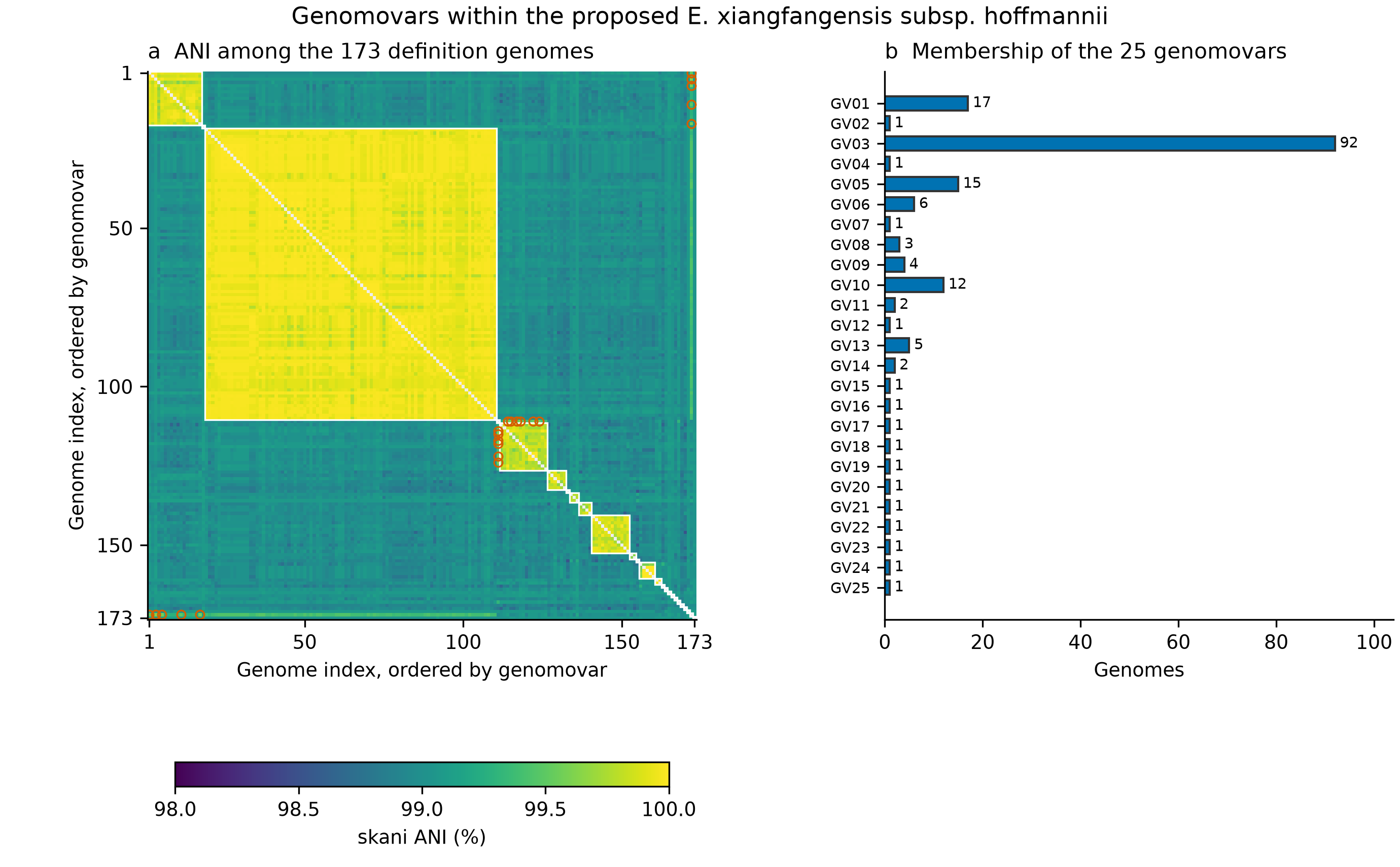

Figure S10. ANI heatmap and sizes of the 25 genomovars; eleven cross-group pairs at or above 99.5% ANI are highlighted.

Panel (a) shows all 14,878 chromosome-scope skani comparisons among the 173 genomes assigned to the proposed *E. xiangfangensis* subsp. *hoffmannii*, ordered by genomovar identifier and then accession. White outlines mark within-genomovar blocks; unmeasured self comparisons are grey. Orange circles mark the eleven distinct pairs from different groups with ANI ≥99.5%, shown in both symmetric positions. All within-group pairs meet the threshold, but some between-group pairs do too. The colour scale is fixed at 98–100%, containing all observed values. Panel (b) gives group sizes, including fifteen singletons. The genome order, all pairwise values and cross-group pairs are supplied with the figure source data; Table S10 provides the membership crosswalk. Pairwise values share genomes and are not independent samples.

##### Figure S11. Pterobacter reference-removal experiments

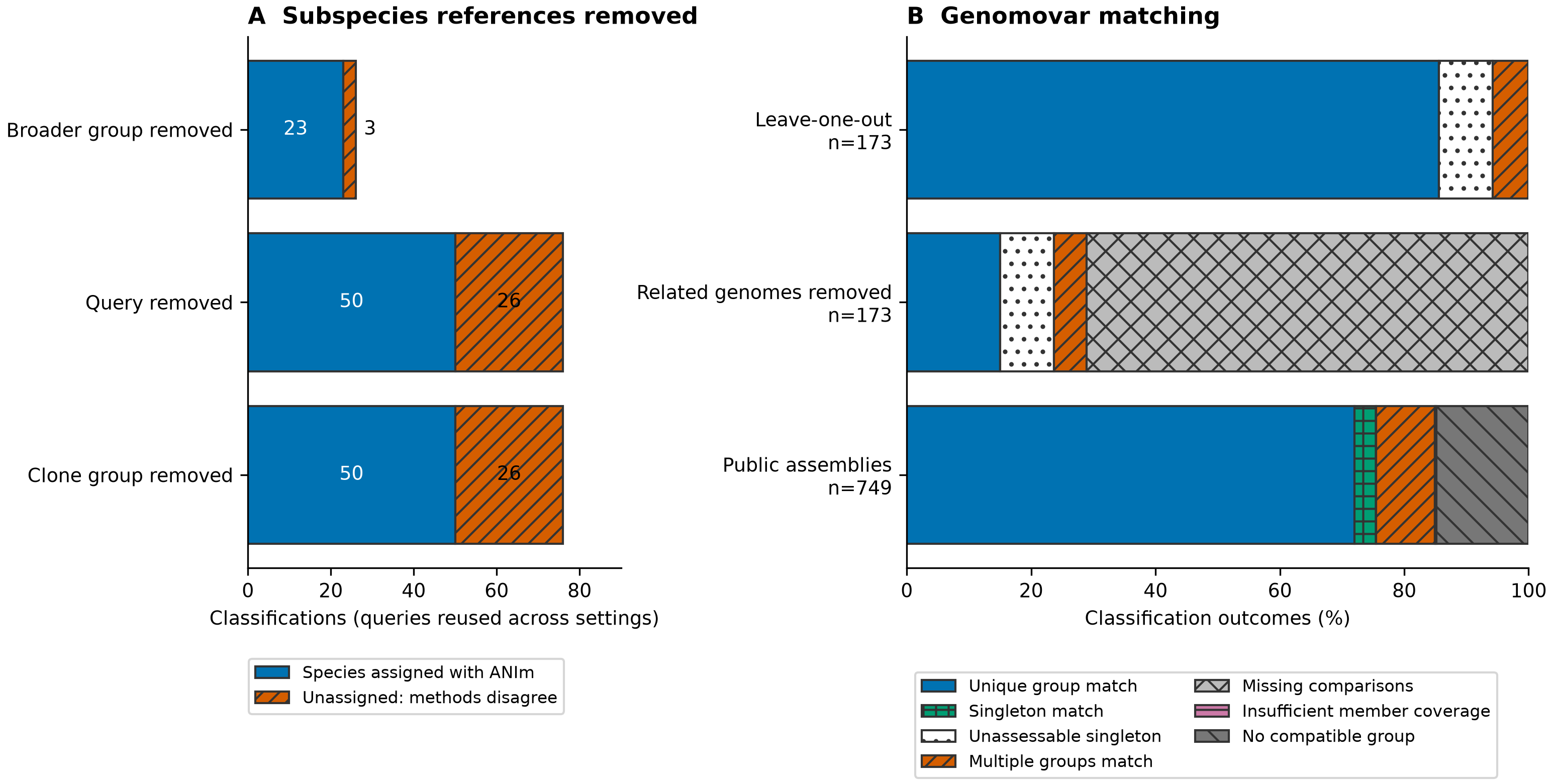

Figure S11. Alignment-assisted species assignments and genomovar matching under reference removal.

(a) Additional species assignments after expected subspecies references were removed, alongside unresolved ANI-method disagreements. The three settings remove the query and aliases, its near-clone block or its broader provenance-linked block; the same queries are reused. (b) Genomovar outcomes with individual queries or related genomes excluded, and for public assemblies assigned to the parent lineage. Each bar has its own denominator. The twelve neighbouring-group matches following complete source-group removal are described separately in S08 and Figure S12.

##### Figure S12. Why omitting a genomovar can create an apparently unique match

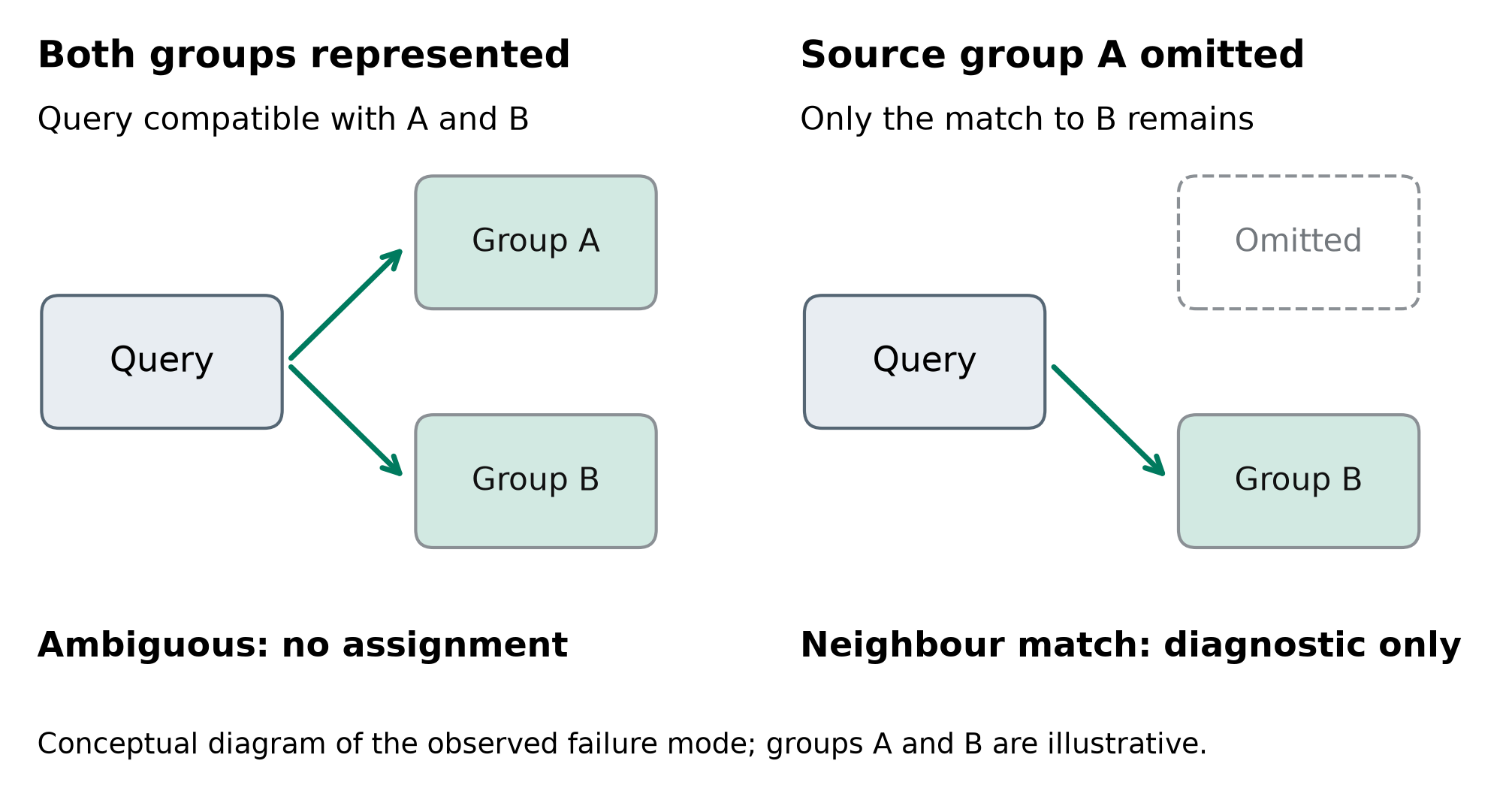

Figure S12. Conceptual diagram of a previously ambiguous query matching a neighbour after its source group is omitted.

With both groups represented, a query compatible with groups A and B has no unique assignment. Removing its source group A leaves only the match to B. Such matches in the reference-removal experiments were recorded as diagnostics with deployment assignments withheld. The letters are illustrative and do not denote the study’s broader taxonomic groups. This schematic explains the twelve observed diagnostic cases; it adds no measurements or biological samples. Eligibility for the parent species and subspecies remains required before genomovar comparison (S08).

##### Figure S13. Kraken names compared with every study-assignment category

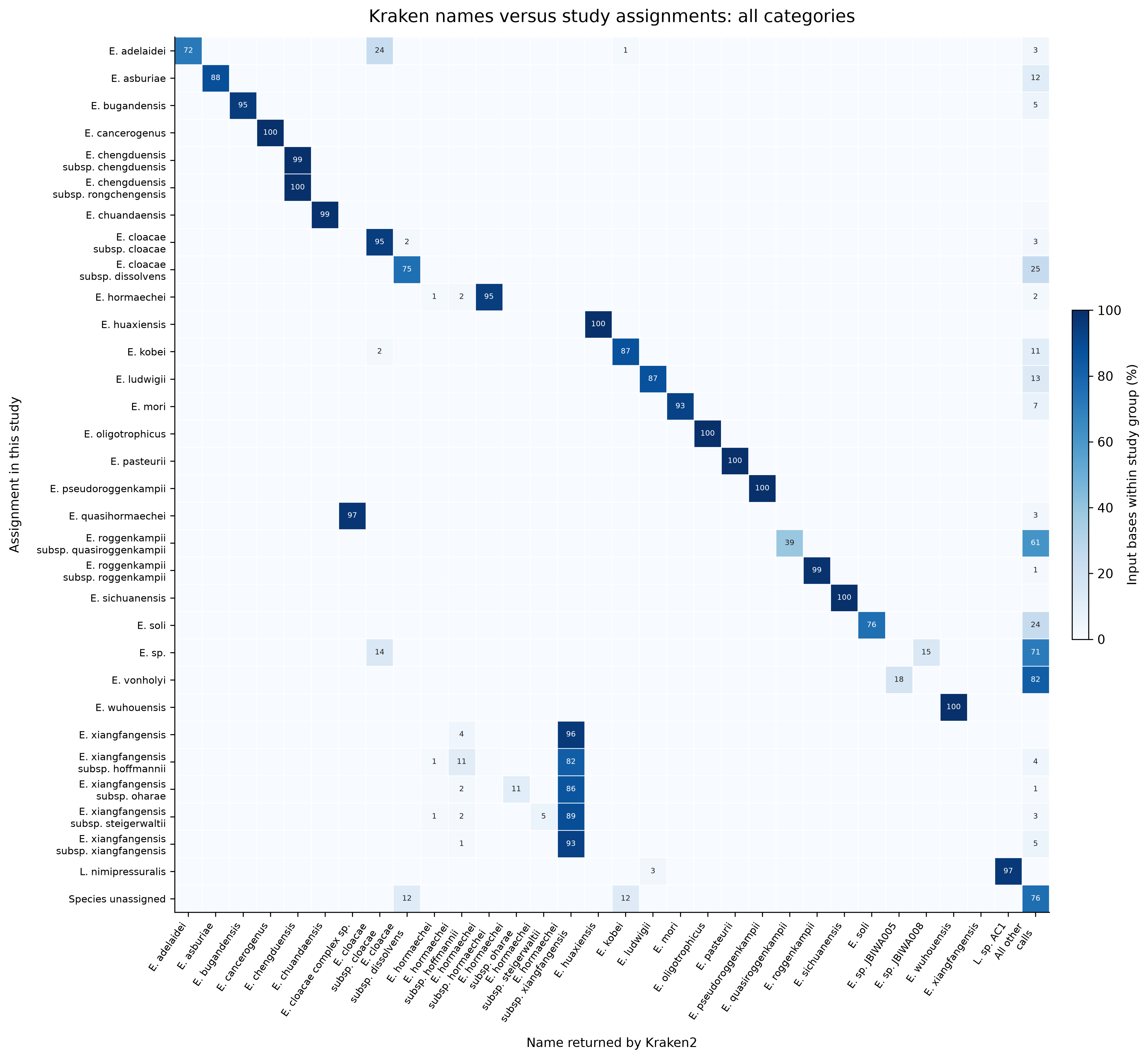

Figure S13. Kraken names versus all 32 revised assignment categories.

The native-contig, default-confidence comparison includes all 32 study-assignment categories, including unnamed and contextual groups. Each cell gives the percentage of that row’s represented input bases carrying the indicated sequence-level call. Strain labels are collapsed to their recorded species or subspecies ancestor. Columns retain the most abundant call in every row and all named columns in Figure 3; all remaining calls are combined in the final column. Numbers are shown for cells at or above 1%, rounded to whole percentages. Exact counts and all 150 ancestor-name columns are retained in the accompanying source table. Different names can refer to compatible taxonomic concepts; this display does not classify every off-diagonal cell as an error.

##### Figure S14. Kraken classification resolution

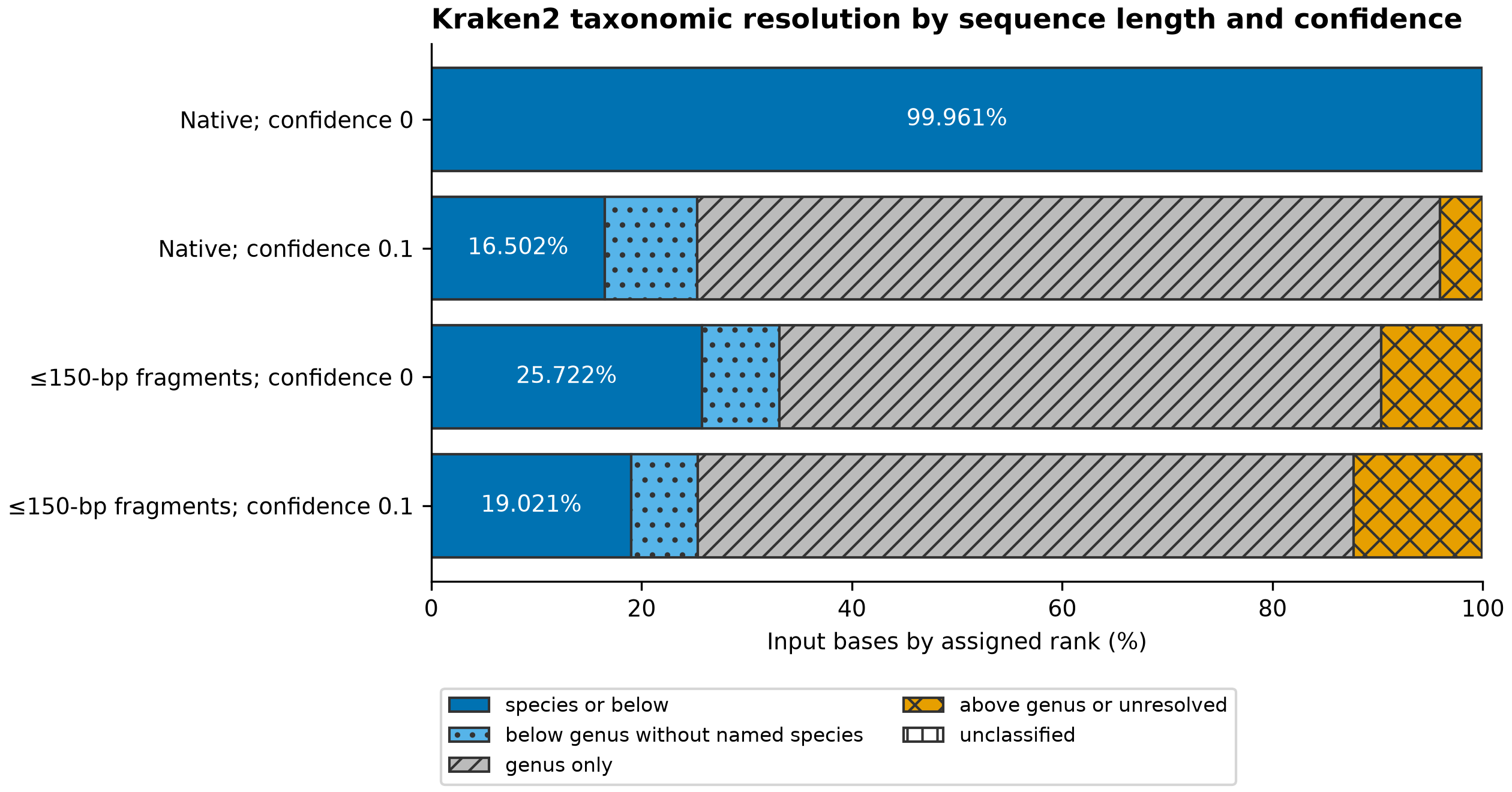

Figure S14. Taxonomic resolution of Kraken calls across sequence lengths and confidence settings.

Native contigs and artificial fragments of at most 150 bp were classified with Kraken2 2.1.6 and Standard 20260226 at confidence 0 and 0.1. Each bar represents the same 5,382,995,503 input bases from 1,070 assemblies, including unclassified sequences. Each call is weighted by the full sequence length.

#### Supplementary tables and source data

The indexed Excel workbook Enterobacter-supplementary-tables.xlsx contains all 23 numbered tables, also supplied as individual TSV files. Accession versions are preserved as text; the new paired ANIm table stores numeric results as numbers.

Tables S01–S23 are supplied as tab-separated files in the supplementary table archive. Each table number identifies one file. The archive also contains structured source-data summaries for dataset overlaps and classifier outcomes. The table numbers below are distinct from the supplement section numbers.

| Table | Contents |
| --- | --- |
| S01 | Assembly accessions, original and revised names, inclusion and quality flags |
| S02 | Dataset selection, exclusions and overlaps |
| S03 | Type-strain figure identifiers, historical names and proposed classifications |
| S04 | Three-method tree comparisons and branch filters |
| S05 | Tested AU hypotheses, results and interpretation limits |
| S06 | Excluded assembly records and chromosome/plasmid similarity categories |
| S07 | Draft-type contig inventory and chromosome/plasmid similarity categories |
| S08 | Nine paired ANIm comparisons before and after removal of candidate plasmid-like contigs |
| S09 | Tree splits in the focused sequence-addition tests |
| S10 | Genomovar membership and definition genomes |
| S11 | Pairs linking different genomovars at or above the ANI threshold |
| S12 | Primary classifier reference panel |
| S13 | Alignment-assisted classification observations by dataset and setting |
| S14 | Comparisons that permitted or prevented alignment-assisted assignment |
| S15 | Twelve matches to neighbouring genomovars after removal of the expected group |
| S16 | Kraken sequence counts, sequence lengths and classification categories |
| S17 | Computational measurements for the 5,803 native queries |
| S18 | Computational measurements for 11,791 reciprocal alignment pairs |
| S19 | Computational measurements for eighteen integrated runs |
| S20 | Proposed name changes and their nomenclatural status |
| S21 | Type-strain culture deposits, assemblies and documentation |
| S22 | Published names and type-genome coverage in the priority check |
| S23 | Genome sizes and G+C contents of the proposed taxa |
